# Hippocampal-cortical connectivity relates to inter-individual differences and training gains in distinguishing similar memories

**DOI:** 10.1101/2025.01.08.631882

**Authors:** P. Iliopoulos, J. Güsten, E. N. Molloy, R. M. Cichy, F. Krohn, A. Maass, E. Düzel

## Abstract

Mnemonic discrimination (MD) is the ability to distinguish current experiences from similar memories. Research on the brain correlates of MD has focused on how regional neural responses are linked to MD. Here we go beyond this approach to investigate inter-regional functional connectivity patterns related to MD, its inter-individual variability and training-related improvement. Based on prior work we focused on medial temporal lobe (MTL), prefrontal cortex (PFC) and visual regions. We used fMRI to determine how functional connectivity patterns between these regions are related to MD before and after 2-weeks of web-based cognitive training. We identified a functional connectivity signature involving MTL-PFC-visual areas during successful MD. We found that hippocampal-PFC connectivity was negatively associated with interindividual variability in MD performance across two different tasks. Hippocampal-PFC connectivity decrease was also linked to interindividual variability in post-training MD improvement. Additionally, training led to increased connectivity from the lateral occipital cortex to the occipital pole area. Our results point to a hippocampal-PFC connectivity pattern which is a reliable, task-invariant, marker of MD performance. This pattern is further related to MD training gains providing causal evidence for its relevance in distinguishing similar memories. Overall, we show that hippocampal-PFC connectivity constitutes a neural resource for MD that enables training-related improvements and could be targeted in future research to enhance cognition.

## 1. Introduction

The ability to remember and distinguish between similar experiences is fundamental to human cognition. This capacity, known as mnemonic discrimination (MD), enables us to form distinct memories. MD is putatively supported by a process called pattern separation, whereby the brain orthogonalizes overlapping neural representations to prevent interference between similar memories (Leal & Yassa, 2018; Yassa & Stark, 2011). MD and pattern separation are predominantly attributed to the hippocampus and associated medial temporal lobe (MTL) structures (Kirwan & Stark, 2007; Leal & Yassa, 2018; Yassa & Stark, 2011). The clinical relevance of MD is illustrated by its decline with aging and Alzheimer’s disease (AD) pathology (Güsten et al., 2021; Leal & Yassa, 2018; Maass et al., 2019; Yassa et al., 2010, 2011). Older adults have intact recognition for exact repetitions of previously encountered stimuli (“repeats”) but they show deficits in identifying similar versions of previously encountered stimuli (“lures”) (Leal & Yassa, 2018; Yassa et al., 2011). This deficit in distinguishing between similar memories is even stronger in individuals with AD, leading to memory confusion and errors (Leal & Yassa, 2018). There is, therefore, a strong interest to understand the neural correlates of MD, the neural basis of its interindividual variability and its improvement by training; not only for elucidating fundamental memory processes but also for developing interventions to mitigate cognitive decline.

In humans, functional magnetic resonance imaging (fMRI) has been instrumental in advancing our knowledge of the neural networks involved in MD. By examining the differential brain activation elicited by similar (lure) versus repeated stimuli—a contrast known as lure detection (LD)—researchers have identified regions implicated in MD and by inference also in pattern separation (Berron et al., 2018). Findings from fMRI studies indicate that in addition to the hippocampus / dentate gyrus (Bakker et al., 2008; Yassa & Stark, 2011), a broader network of brain regions are implicated in MD (Amer & Davachi, 2023; Pidgeon & Morcom, 2016). These include other MTL regions (Reagh & Yassa, 2014), the prefrontal cortex (PFC), and areas in the visual cortex (Amer & Davachi, 2023; Klippenstein et al., 2020; Nash et al., 2021; Pidgeon & Morcom, 2016; Wais et al., 2017).

Here we addressed the question how MTL, PFC and visual areas functionally interact to support MD. To that end, we took a two-pronged approach. First, we determined how functional connectivity between these regions during an MD task related to inter-individual variability in performance across two different MD paradigms. Second, we performed a MD training intervention and determined whether training-related gains in MD performance were related to the same functional connectivity patterns that also explained interindividual variability. Our rationale is that the convergence between functional connectivity related to interindividual variability and training-related gains can uncover neural resources that underlie MD more robustly than the cross-sectional and univariate studies up to date.

Our study builds on previous work showing that it is possible to enhance MD performance with cognitive training. A recent study performed in our lab reported decreased false recognition of lure stimuli and regional brain activation changes following a 2-week web-based cognitive intervention (Güsten et al., 2024). Here we determined whether cognitive training altered functional connectivity within the MD network and whether these changes resembled the patterns displayed in high-performing individuals. We also tested whether such a connectivity pattern relates to performance in another MD task (Stark et al., 2019). An overarching question is whether neural resources explaining interindividual variability also explain training-related gains. If we find convergence -the same neural networks are involved in both individual differences and training gains-it would indicate that high performers and beneficiaries of cognitive training tap into similar neural resources. This would indicate that these networks are fundamental to MD and strengthen our confidence in targeting them in future research. Conversely, if there is no convergence, it would imply that training induces neural adaptations distinct from the observed inter-individual variability patterns. Understanding whether there is convergence contributes to elucidating the neural correlates of MD and developing effective interventions.

We hypothesize that effective MD is supported by enhanced bidirectional connectivity among a MTL-PFC-visual network. Our rationale is that such connectivity would facilitate the integration of visual information with memory processes. We expect specifically hippocampal-PFC interactions to play a central role, given findings indicating a role of PFC, alongside the hippocampus, in MD both in humans (Amer & Davachi, 2023; Nash et al., 2021; Pidgeon & Morcom, 2016; Wais et al., 2017, 2018) and animal studies (Wang et al., 2021). We further hypothesize that training will lead to altered connectivity within the MD-associated network, reflecting neural plasticity in the form of functional reorganization (A. M. C. Kelly & Garavan, 2005; C. Kelly & Castellanos, 2014). We expect the connectivity changes to converge with the connectivity pattern related to inter-individual performance differences.

For these objectives, we employed a two-week web-based cognitive training protocol emphasizing MD, using a 2×2 longitudinal intervention design (Group × Time) (Güsten et al., 2024). Functional connectivity was assessed through region-of-interest generalized psychophysiological interaction (gPPI) analyses (McLaren et al., 2012), allowing us to model event-related effective connectivity between brain regions while accounting for task-related activity. Participants underwent pre- and post-training fMRI scanning at 3 Tesla performing an established MD task, which includes both object and scene stimuli (Berron et al., 2018; Güsten et al., 2021). Our primary outcome measure was the functional connectivity associated with correct lure detection compared to repetition trials. Additionally, the participants completed a battery of behavioral paradigms that included another established MD task, the mnemonic similarity task (MST) (Stark et al., 2019). We hypothesize that the connectivity pattern related to inter-individual differences will be linked to performance measured in this different MD task.

By integrating analyses of functional connectivity with cognitive training, our study aims to advance the understanding of the neural mechanisms underlying MD and their potential plasticity. This knowledge could inform the development of targeted interventions to enhance memory function and contribute to preventative strategies against cognitive decline in aging and neurodegenerative diseases.

## 2. Results

To assess the impact of cognitive training on mnemonic discrimination (MD) and its neural mechanisms, we analyzed behavioral and fMRI data from 54 healthy young adults who completed pre- and post-intervention sessions. Participants were randomly assigned to either the experimental training group (n=26), which received MD-focused cognitive training, or the active control group (n=27), which engaged in a psychomotor task without an MD component (Fig.1). During fMRI scanning sessions, all participants performed a 6-back object-scene recognition task that required distinguishing between highly similar (“lure”) and identical (“old”) images, engaging MD (Fig.1A-C). Detailed descriptions of the participant groups, behavioral tasks and training protocol are provided in the Methods section and in (Güsten et al., 2024). For an overview of the analysis flow see Fig. 2.

**Figure 1.**
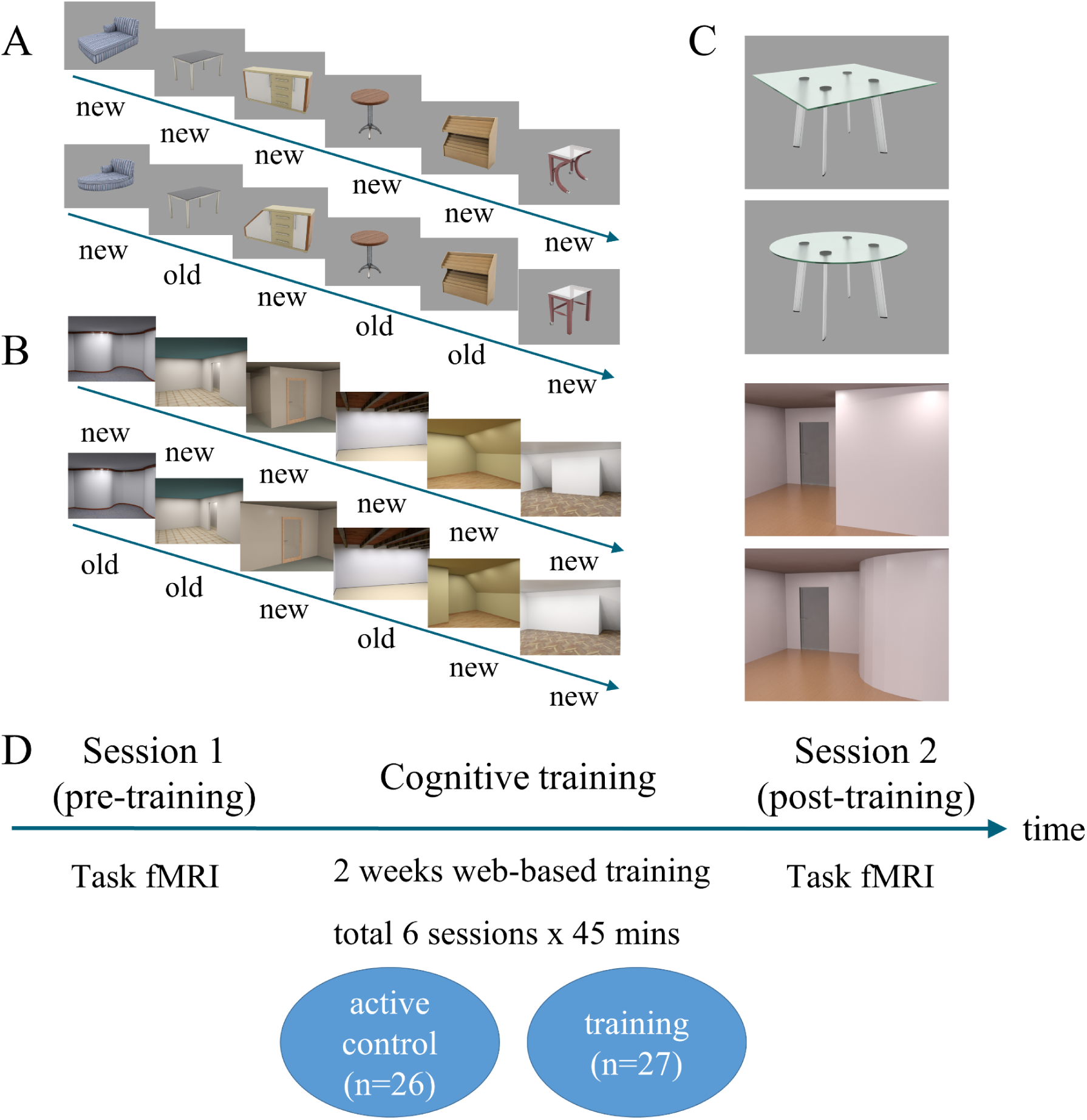
Experimental 6-back object-scene mnemonic discrimination paradigm and study design. **A-C**: The 6-back object-scene mnemonic discrimination (MD) task was performed in the scanner (fMRI) and remotely on a computerized web-based training platform. Stimuli were presented in sequences of 12 items: 6 encoding (‘first’ trials) followed by 6 test phase images. Participants were asked to respond “new” or “old” to each image using their middle and index fingers. In the test phase, they had to recognize whether the stimulus was similar but slightly changed (**‘**lure**’** correct response: ‘new’) or an identical repetition (**‘**repeat**’** correct response: ‘old’) compared to the ‘first’ images. Each sequence consisted of either objects (A) or scenes (B) only. Lure and repeat stimuli only differed in shape or geometry (C). **D**. Participants completed the 6-back object-scene task inside the MRI scanner. They were scanned pre- and post a 2-week remote web-based cognitive training intervention that consisted of three 45-minute sessions per week. One group (n=26) trained using a web-based version of the above task, whereas an active control group (n=27) was presented with the same images but did a psychomotor task by clicking on moving neurons-icons on top of the images. Note that different sets of images were shown for each fMRI session, and the cognitive training.

**Figure 2.**
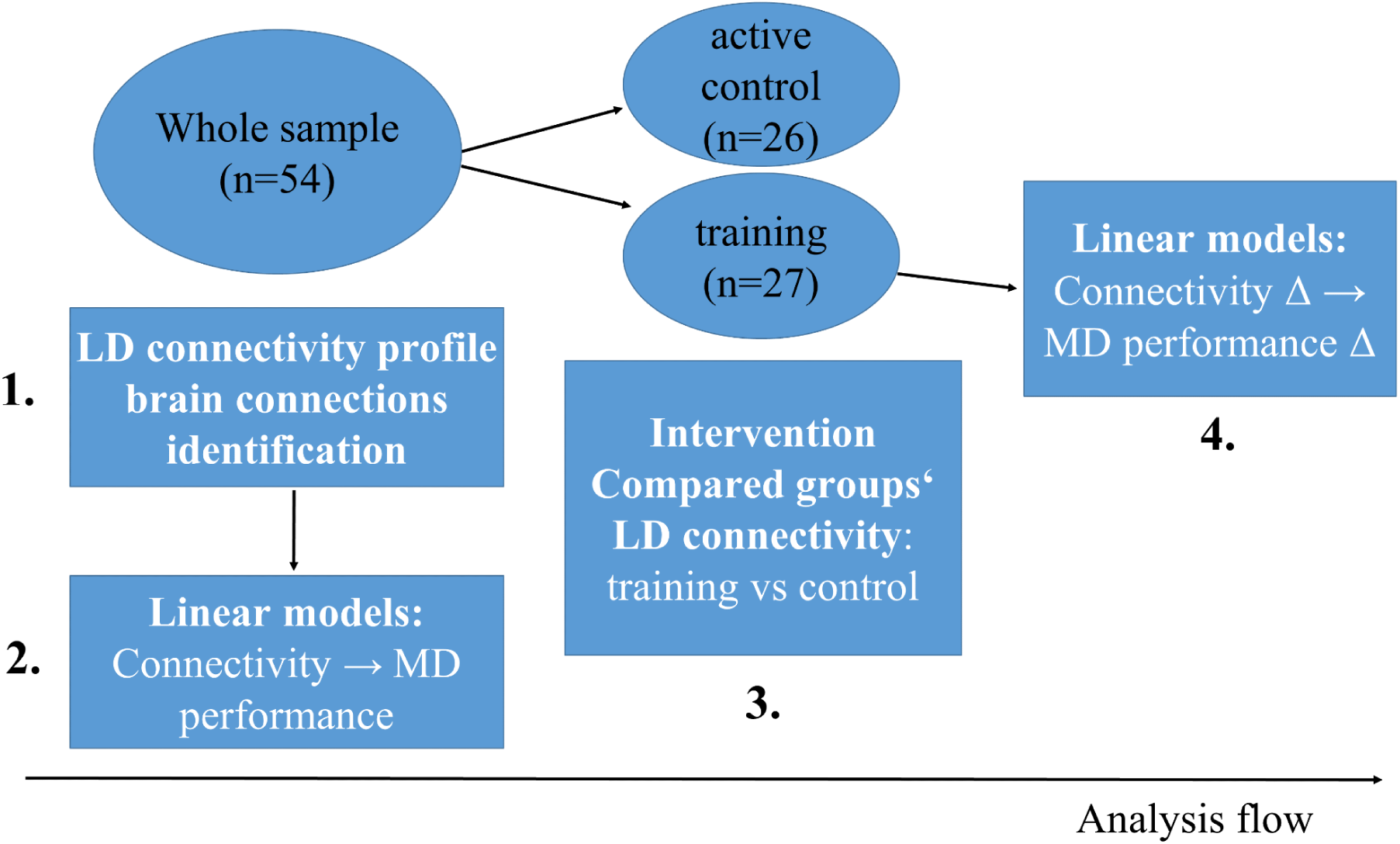
Study analysis flow. **1)** To identify the lure detection (LD) connectivity profile (i.e. significant brain connections), we first performed connectivity analyses in the whole sample pre-training data (n=54). **2)** Then, focusing on the identified brain connections from the step 1, we fitted linear models to test the link between connectivity in each connection and MD performance at the baseline whole sample data. **3)** Next, we compared the LD connectivity for the training versus control group for each connection identified in step 1. Note: One dataset couldnt be analyzed here due to technical error. **4)** Then, we fitted linear models to test the link between connectivity change and MD performance change (Δ: post-training versus pre-training change); we focused on the brain connections which exhibited a significant link to MD performance at baseline or a training effect (i.e. the significant connections from steps 2 and 3 respectively).

### 2.1 Lure detection connectivity pattern involves an MTL-PFC-visual network

To first characterize the connectivity pattern during successful MD, we contrasted the event-related connectivity for the correct lure versus repeat trials. We term this difference the lure detection contrast (LD) and it is used for all connectivity analyses in our study. We performed gPPI ROI-to-ROI analysis in the pre-training whole-sample data, using cluster-based inference and an a-priori selection of ROIs that included major prefrontal (PFC), medial temporal lobe (MTL) and visual areas given our a-priori interest in the role of MTL-PFC-visual connections in MD.

In accordance with our hypothesis, we observed three significant clusters (p < .05 p-FDR) (see Fig. 3; Table 1 for statistics) for the LD contrast. The first cluster displayed reduced connectivity in visual-to-visual and visual-MTL connections during correct lure detection, specifically between the lateral occipital cortex (LOC) and occipital pole (OP), and between the OP and perirhinal (PRC) as well as OP and entorhinal cortex (EC). The second cluster exhibited higher LD connectivity in LOC – PFC and inferior frontal gyrus pars triangularis (IFG tri)-MTL connections: specifically, between the LOC and the IFG tri, superior frontal gyrus (SFG), and inferior frontal gyrus pars opercularis (IFG oper) areas. In addition, we found higher LD connectivity between the LOC and the hippocampus, and between the IFG tri and the EC as well as PRC. The third cluster exhibited higher connectivity during correct lure detection in mainly hippocampal-PFC connections: specifically between the hippocampus and the SFG, IFG oper and IFG tri areas. In summary, we found that MD is associated with lower task-based connectivity in LOC-OP and OP – MTL connections, and higher connectivity in LOC – PFC, LOC-hippocampus, IFG tri-MTL and hippocampal-PFC connections. These clusters reveal a task connectivity profile associated with successful MD that involve specific MTL-PFC-visual connections. Thus, all our subsequent connectivity analyses focused on these identified MD-associated functional networks (Figure 3A).

**Figure 3.**
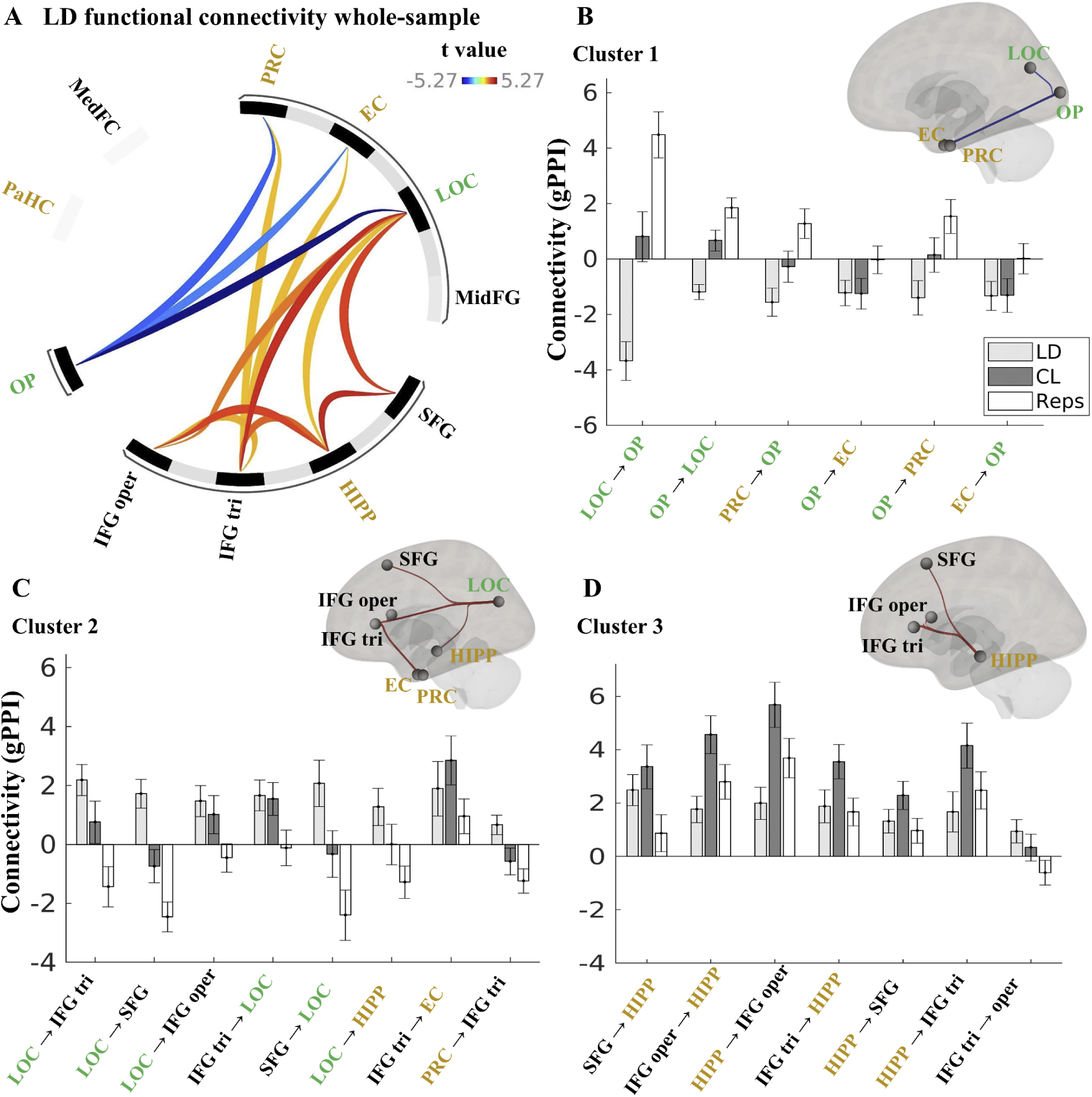
A. Task connectivity profile during successful MD. LD contrast gPPI values: ie. event-related connectivity in correct lures (CL) versus repeats (Reps) trials. Three significant clusters were observed for the LD contrast : 1) lower connectivity for LOC – OP and OP - MTL (EC, PRC) connections; 2) higher connectivity for LOC – PFC areas and IFG tri to LOC, EC, PRC; 3) higher connectivity for hippocampal - PFC connections (see Table 1 for more detail). Color represents the t values. **B-D**. Barplots for each cluster 1-3 plotted separately. Y axis shows the mean gPPI value for each of the three conditions (ie: LD contrast; correct lures and repeats minus the perceptual control trials). **PaHC**: parahippocampal cortex, **MedFC**: medial frontal cortex, **PRC**: perirhinal cortex, **EC**: enthorhinal cortex, **LOC**: lateral occipital cortex, **MidFG**: middle frontal gyrus, **SFG**: superior frontal gyrus, **HIPP**: hippocampus, **IFG**: inferior frontal gyrus; **tri**: pars triangularis, **oper**: pars opercularis. **OP**: occipital pole. The colour of the ROIs’ names denote whether they are a visual (green), MTL (gold), or PFC (black) area.

**Table 1.** ROI-to-ROI connectivity (gPPI) for the lure detection (LD) contrast.

| Cluster/Connection | Statistic | *p* | *p-FDR* |
| --- | --- | --- | --- |
| *Cluster 1* | $F(2,52) = 15.15$ | $< .001$ | $< .001$ |
| LOC - OP | $T(53) = -5.27$ | $< .001$ | $< .001$ |
| OP - LOC | $T(53) = -4.36$ | $< .001$ | 0.001 |
| PRC - OP | $T(53) = -3.05$ | 0.004 | 0.036 |
| OP - EC | $T(53) = -2.67$ | 0.010 | 0.051 |
| OP - PRC | $T(53) = -2.24$ | 0.029 | 0.098 |
| EC - OP | $T(53) = -2.54$ | 0.029 | 0.098 |
| *Cluster 2* | $F(2,52) = 8.16$ | 0.001 | 0.003 |
| LOC - IFG tri | $T(53) = 4.17$ | $< .001$ | 0.001 |
| LOC - SFG | $T(53) = 3.55$ | 0.001 | 0.003 |
| LOC - IFG oper | $T(53) = 2.80$ | 0.007 | 0.014 |
| IFG tri - LOC | $T(53) = 3.21$ | 0.002 | 0.019 |
| SFG - LOC | $T(53) = 2.65$ | 0.002 | 0.019 |
| LOC - HIPP | $T(53) = 2.03$ | 0.048 | 0.080 |
| IFG tri - EC | $T(53) = 2.05$ | 0.046 | 0.106 |
| PRC - IFG tri | $T(53) = 2.01$ | 0.050 | 0.152 |
| *Cluster 3* | $F(2,52) = 8.14$ | 0.001 | 0.003 |
| SFG - HIPP | $T(53) = 4.28$ | $< .001$ | 0.001 |
| IFG oper - HIPP | $T(53) = 3.56$ | 0.001 | 0.008 |
| HIPP - IFG oper | $T(53) = 3.29$ | 0.002 | 0.018 |
| IFG tri - HIPP | $T(53) = 3.04$ | 0.004 | 0.019 |
| HIPP - SFG | $T(53) = 2.96$ | 0.005 | 0.023 |
| HIPP - IFG tri | $T(53) = 2.22$ | 0.031 | 0.103 |
| IFG tri - IFG oper | $T(53) = 2.17$ | 0.035 | 0.106 |

*Note:* Parametric multivariate statistics (cluster-based inference, functional network connectivity). Cluster-level *p*-FDR corrected (MVPA omnibus test), connection threshold: $p < .05$ *p*-uncorrected. **PRC**: perirhinal cortex, **EC**: entorhinal cortex, **HIPP**: hippocampus, **LOC**: lateral occipital cortex, **OP**: occipital pole, **SFG**: superior frontal gyrus, **IFG**: inferior frontal gyrus, **tri**: triangularis, **oper**: opercularis.

#### 2.2.1 Higher LD connectivity in hippocampal-PFC connections is linked to poorer mnemonic discrimination performance

Following the characterization of baseline MD-associated functional connectivity, we investigated whether there is a quantitative relationship between this LD task connectivity in each cluster and MD behavioral performance (defined using the *A* discriminability index). To test this, we fitted linear regression models between the LD connectivity in each connection and the *A* discriminability index (covariates: age, sex).

Results show that lower LD task connectivity in hippocampal-PFC connections (cluster 3) is associated with higher MD (see Fig.4, Table 2). The following connections exhibited a significant negative relationship: SFG-hippocampus, hippocampus-IFG oper, IFG tri-hippocampus, hippocampus-SFG, and hippocampus-IFG tri (see Table 2 for statistics).

**Figure 4.**
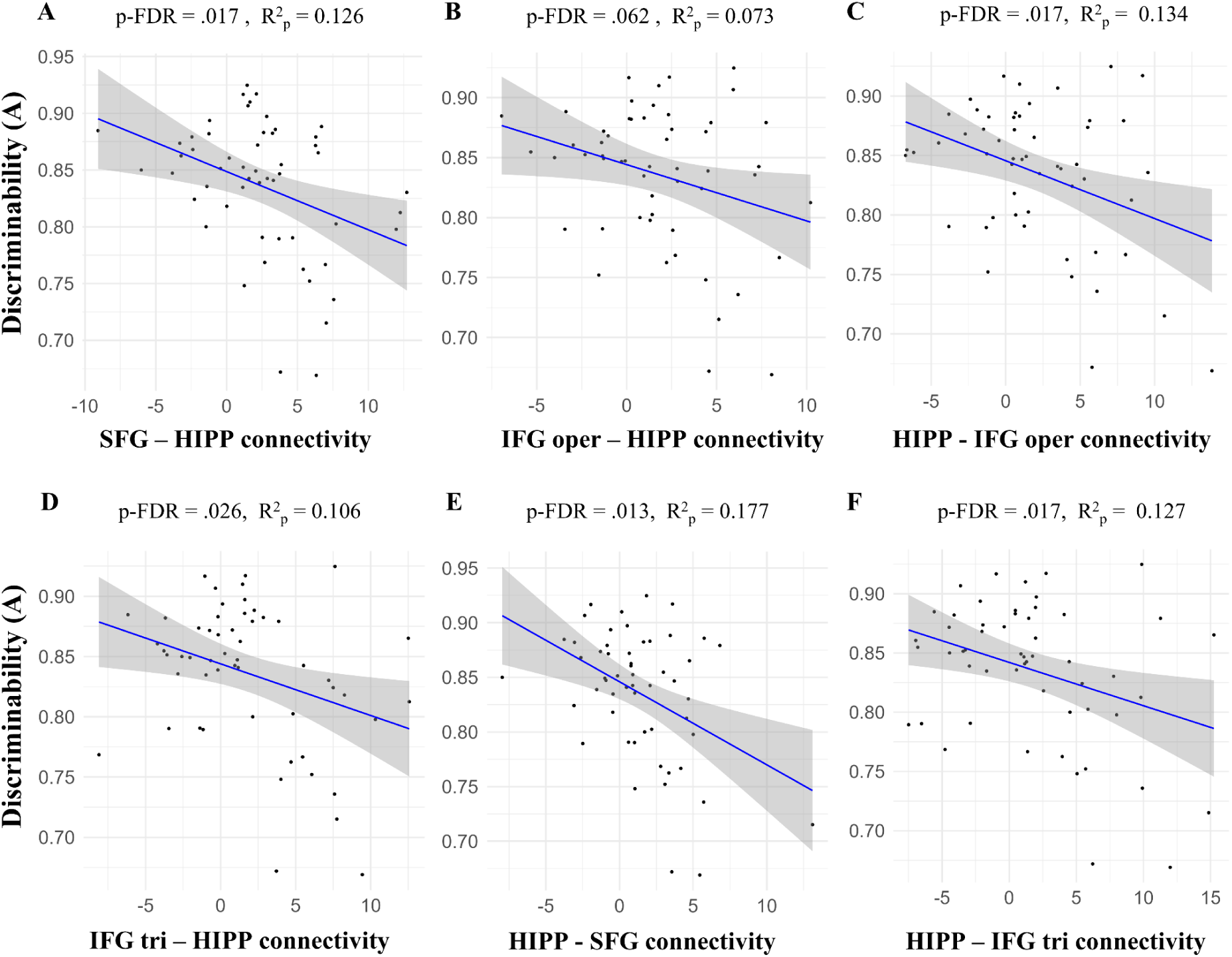
Brain connectivity-behavior linear regression: hippocampal-PFC connections. The X axis depicts the independent variable: LD connectivity values (gPPI) in a certain connection (seed ROI – target ROI). The Y axis shows the dependent variable: A discriminability. The p-FDR corrected and partial R square values are shown. FDR-correction was applied to all the connections within this cluster. In all connections depicted (A-F) we find a negative link with lower connectivity being associated with higher memory performance.

**Table 2.** Linear model regression results for the effect of connectivity on MD performance (A).

| Cluster | Connection | b | SE | *p* | *p*-FDR | R<sup>2</sup><sub>p</sub> |
| --- | --- | --- | --- | --- | --- | --- |
| 1 | LOC-OP | 0.000 | 0.002 | 0.780 | 0.780 | 0.002 |
|  | OP-LOC | 0.008 | 0.004 | 0.055 | 0.164 | 0.072 |
|  | PRC-OP | -0.003 | 0.002 | 0.130 | 0.260 | 0.045 |
|  | OP-EC | -0.002 | 0.002 | 0.469 | 0.586 | 0.011 |
|  | OP-PRC | -0.001 | 0.002 | 0.489 | 0.586 | 0.010 |
|  | EC-OP | -0.004 | 0.002 | **0.047** | 0.164 | 0.077 |
| 2 | LOC-IFG tri | -0.002 | 0.002 | 0.305 | 0.526 | 0.048 |
|  | LOC-SFG | -0.004 | 0.002 | 0.129 | 0.444 | 0.045 |
|  | LOC-IFG oper | -0.003 | 0.002 | 0.133 | 0.444 | 0.045 |
|  | IFG tri-LOC | -0.001 | 0.002 | 0.533 | 0.609 | 0.008 |
|  | SFG-LOC | 0.001 | 0.001 | 0.679 | 0.679 | 0.003 |
|  | LOC-HIPP | -0.002 | 0.002 | 0.329 | 0.526 | 0.019 |
|  | IFG tri-EC | -0.002 | 0.001 | 0.166 | 0.444 | 0.038 |
|  | PRC-IFG tri | 0.002 | 0.003 | 0.513 | 0.609 | 0.009 |
| 3 | SFG-HIPP | -0.005 | 0.002 | **0.010** | **0.017** | 0.126 |
|  | IFG oper-HIPP | -0.004 | 0.002 | 0.053 | 0.062 | 0.073 |
|  | HIPP-IFG oper | -0.005 | 0.002 | **0.008** | **0.017** | 0.134 |
|  | IFG tri-HIPP | -0.004 | 0.002 | **0.019** | **0.026** | 0.106 |
|  | HIPP-SFG | -0.008 | 0.002 | **0.002** | **0.013** | 0.177 |
|  | HIPP-IFG tri | -0.004 | 0.001 | **0.010** | **0.017** | 0.127 |
|  | IFG tri-IFG oper | -0.001 | 0.003 | 0.698 | 0.698 | 0.003 |

*Note.* A model regressing connectivity on the outcome measure A (age, sex: covariates of no interest) was fitted for each brain connection within the 3 clusters. Given are the b=beta estimates, SE=standard error, p-values. Within cluster p-FDR correction was applied to the p-values. R<sup>2</sup><sub>p</sub> = partial R square is given to estimate the variance explained by each connection independently of the covariates (age, sex). p-FDR < 0.05 are considered as significant. **PRC**: perirhinal cortex, **EC**: entorhinal cortex, **HIPP**: hippocampus, **LOC**: lateral occipital cortex, **OP**: occipital pole, **SFG**: superior frontal gyrus, **IFG**: inferior frontal gyrus, **tri**: triangularis, **oper**: opercularis.

Conversely, we observed no significant relationship between LD connectivity and MD for any connection within clusters 1 and 2. Within cluster 1, there was a non-significant positive association between OP-LOC connectivity and MD performance (A index) (*p* = .055, *p-FDR* = .164, R^2^_p_ = 0.077), and a negative link between EC-OP connectivity and MD performance that did not survive correction (*p* = .047, *p-FDR* = .164, R^2^_p_ = 0.077) (Table 2).

In sum, lower task-based connectivity during MD between the hippocampus and PFC areas including the SFG, IFG oper and IFG tri is associated with better MD performance, as measured with the *A discriminability index*.

#### 2.2.2 Higher connectivity in the hippocampal-PFC network is linked with poorer performance in out-of the scanner memory task measures

To examine the link between the LD task connectivity in the hippocampal-PFC network (Cluster 3) and performance in various independent memory tasks, we conducted a multivariate regression analysis. This analysis used the LD hippocampal-PFC composite connectivity score as a predictor and a series of memory task measures as dependent variables (see Table 4 for details). The results revealed a significant multivariate effect (Pillai = 0.351, F(6, 36) = 3.25, p = .012), suggesting that variations in hippocampal-PFC connectivity significantly influence overall performance on these tasks (Table 3). Specifically, higher connectivity scores were associated with poorer performance on the lure discrimination index (LDI) and the corrected hit rate of the mnemonic similarity task (MST; mnemonic discrimination task), highlighting a targeted impact (Table 4). Conversely, no significant effects were observed for the other memory measures examined. This pattern underscores the specificity of connectivity effects to particular cognitive processes involved in MD.

**Table 3.** Multivariate regression model of the composite connectivity score (hippocampal-PFC) effect on the combined behavioral measures.

| IV | Pillai test | F | DF<sub>n,d</sub> | p |
| --- | --- | --- | --- | --- |
| Composite Connectivity Score | 0.351 | 3.25 | 6, 36 | **0.012** |
| Age | 0.193 | 1.44 | 6, 36 | 0.228 |
| Sex | 0.175 | 1.27 | 6, 36 | 0.294 |

*Note.* Multivariate regression model. The LD composite connectivity score in the hippocampal-PFC connections is the main predictor variable (age, sex: covariates of no interest). The combined behavioral measures (described in Table 4) are the dependent variables. IV: independent variable.

**Table 4.**
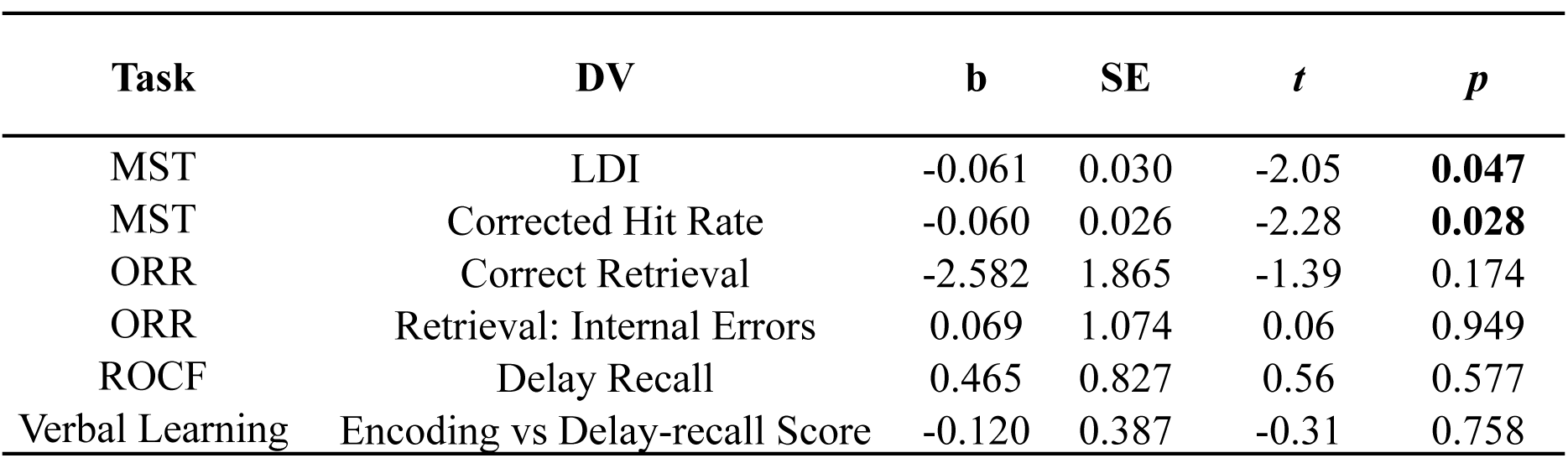
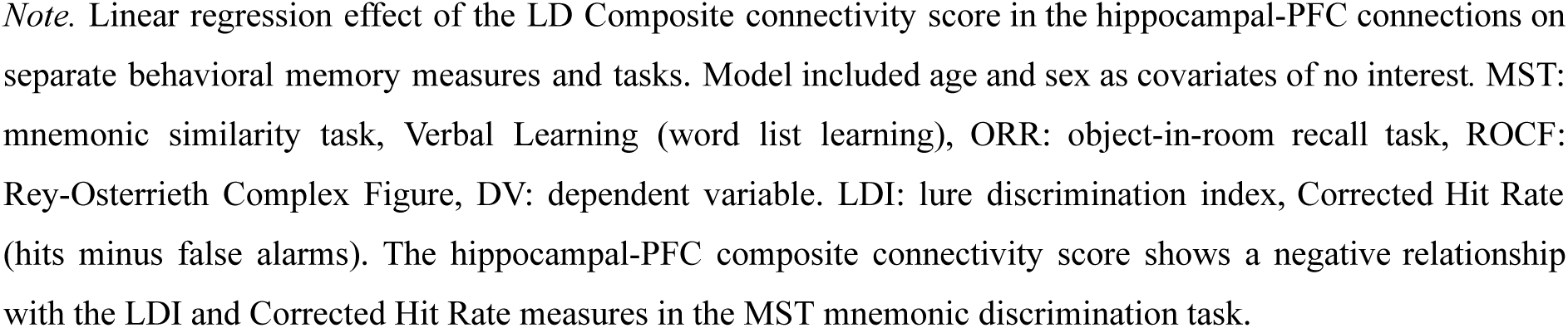
Effects of composite connectivity score (hippocampal-PFC) on memory behavioral measures.

| Task | DV | b | SE | t | p |
| --- | --- | --- | --- | --- | --- |
| MST | LDI | -0.061 | 0.030 | -2.05 | **0.047** |
| MST | Corrected Hit Rate | -0.060 | 0.026 | -2.28 | **0.028** |
| ORR | Correct Retrieval | -2.582 | 1.865 | -1.39 | 0.174 |
| ORR | Retrieval: Internal Errors | 0.069 | 1.074 | 0.06 | 0.949 |
| ROCF | Delay Recall | 0.465 | 0.827 | 0.56 | 0.577 |
| Verbal Learning | Encoding vs Delay-recall Score | -0.120 | 0.387 | -0.31 | 0.758 |

*Note.* Linear regression effect of the LD Composite connectivity score in the hippocampal-PFC connections on separate behavioral memory measures and tasks. Model included age and sex as covariates of no interest. MST: mnemonic similarity task, Verbal Learning (word list learning), ORR: object-in-room recall task, ROCF: Rey-Osterrieth Complex Figure, DV: dependent variable. LDI: lure discrimination index, Corrected Hit Rate (hits minus false alarms). The hippocampal-PFC composite connectivity score shows a negative relationship with the LDI and Corrected Hit Rate measures in the MST mnemonic discrimination task.

### 2.3 Cognitive training effects

#### 2.3.1 Improved mnemonic discrimination performance after training

The behavioral results from this study were previously reported in Güsten et al. (2024). The 2-week training led to reduced false alarms (incorrect lure) without affecting hits, suggesting improved performance on correct lures. Here, we further analyzed the bias-corrected A discriminability index. After the baseline scan, where all participants were assessed as one group, they were divided into a training group (n = 27) and an active control group (n = 26).

A 2-way mixed ANOVA (covariates: age, sex) testing the Group (training, control) × Time (pre-, post-training) interaction on the A discriminability index revealed a significant effect (F(1,49) = 9.34, p = 0.004, η²G = .045) (Fig. 5, Table 5). Simple effects analyses showed no pre-training difference between the training group (M = 0.842, SE = 0.012) and the control group (M = 0.836, SE = 0.011) [t(49) = 0.361, p = 0.719]. Post-training, the training group (M = 0.917, SE = 0.009) outperformed the control group (M = 0.867, SE = 0.009) [t(49) = 4.018, p < 0.001] (Fig. 5A, Table 5). These results show a training-induced improvement in A discriminability index, driven by increased correct lure responses (see Fig. 5).

**Figure 5.**
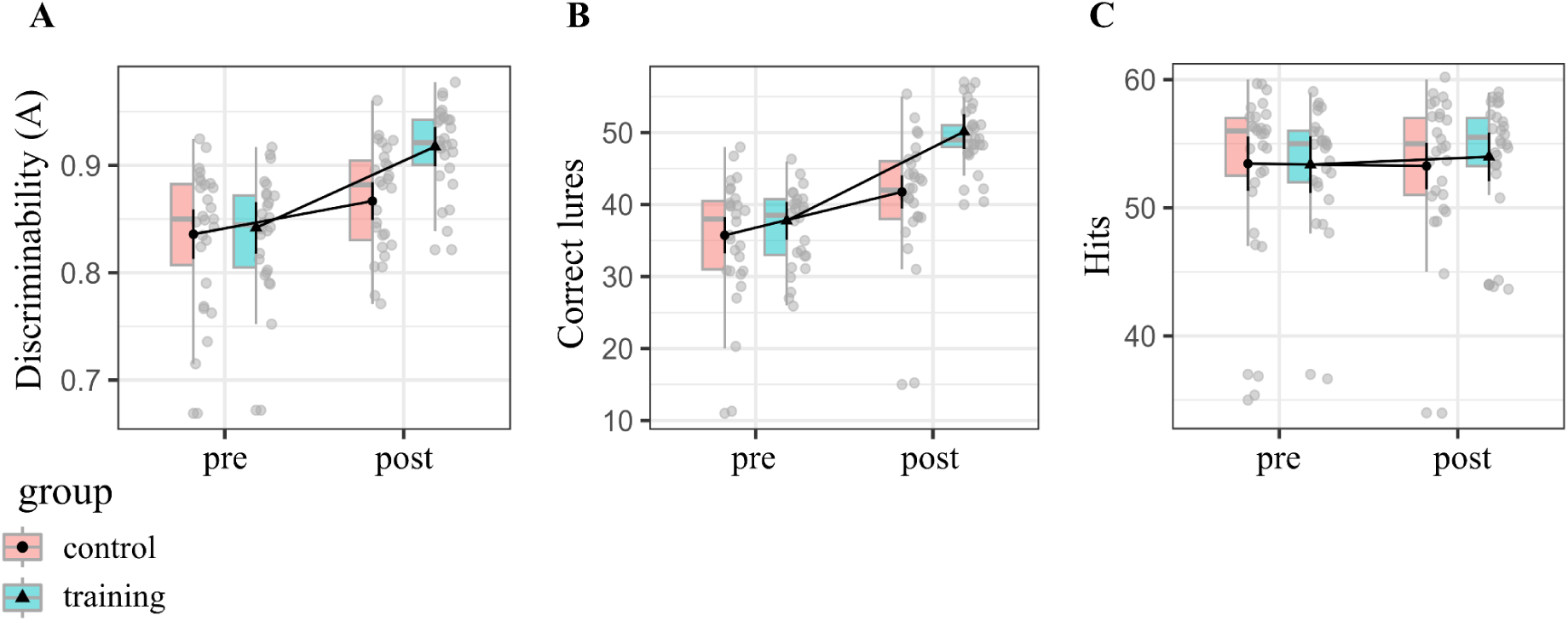
Behavioral training effects. Depicted is the Group x Time (pre-, post-training) interaction shown for different measures: A) for the discriminability index A; B) lure correct responses; C) correct repeat trials (“Hits”). We found a significant Group x Time interaction for the A Discriminability, which is driven by correct rejections (see also Güsten et al. 2024).

**Table 5.** ANOVA results for discriminability index A.

| DV | Effect | Condition | Mean | SE | DF<sub>n,d</sub> | F | p | $\eta^2_G$ | $\eta^2_p$ |
| --- | --- | --- | --- | --- | --- | --- | --- | --- | --- |
| A | Group | Control | 0.851 | 0.009 | 1,49 | 4.94 | 0.031 | 0.070 | 0.092 |
|  |  | Training | 0.880 | 0.009 |  |  |  |  |  |
|  | Time | Pre | 0.839 | 0.008 | 1,49 | 52.59 | < .001 | 0.210 | 0.518 |
|  |  | Post | 0.892 | 0.006 |  |  |  |  |  |
|  | Group x Time | Control / Pre | 0.836 | 0.011 | 1,49 | 9.34 | 0.004 | 0.045 | 0.160 |
|  |  | Training / Pre | 0.842 | 0.012 |  |  |  |  |  |
|  |  | Control / Post | 0.867 | 0.009 |  |  |  |  |  |
|  |  | Training / Post | 0.917 | 0.009 |  |  |  |  |  |

*Note.* Cognitive training results using ANOVA on A discriminability values with factors “Group” (control, training; between-subject) and Time (pre, post-training; within-subject). Group x Time interaction and main effects are shown. Given are mean values and standard error of the mean (SE), degrees of freedom for the numerator (DFn) and denominator (DFd), p-values (p), and effect size ( $\eta^2_g, \eta^2_p$ ). P-values < 0.05 are considered as significant. DV= dependent variable.

#### 2.3.2. Increased intra-occipital cortex connectivity after cognitive training

Next we investigated the effects of cognitive training on the LD contrast. To assess training-induced effects within each of the three clusters identified above (see Figure 6A-C), we used mixed ANOVAs (Group x Time interaction) with the LD contrast metrics as the dependent variable (covariates: age, sex). Via these interactions, we tested whether the connectivity change in each connection was different for the training versus the control group. For detailed statistics for each connection/cluster refer to supplementary Tables S1-S3.

**Figure 6.**
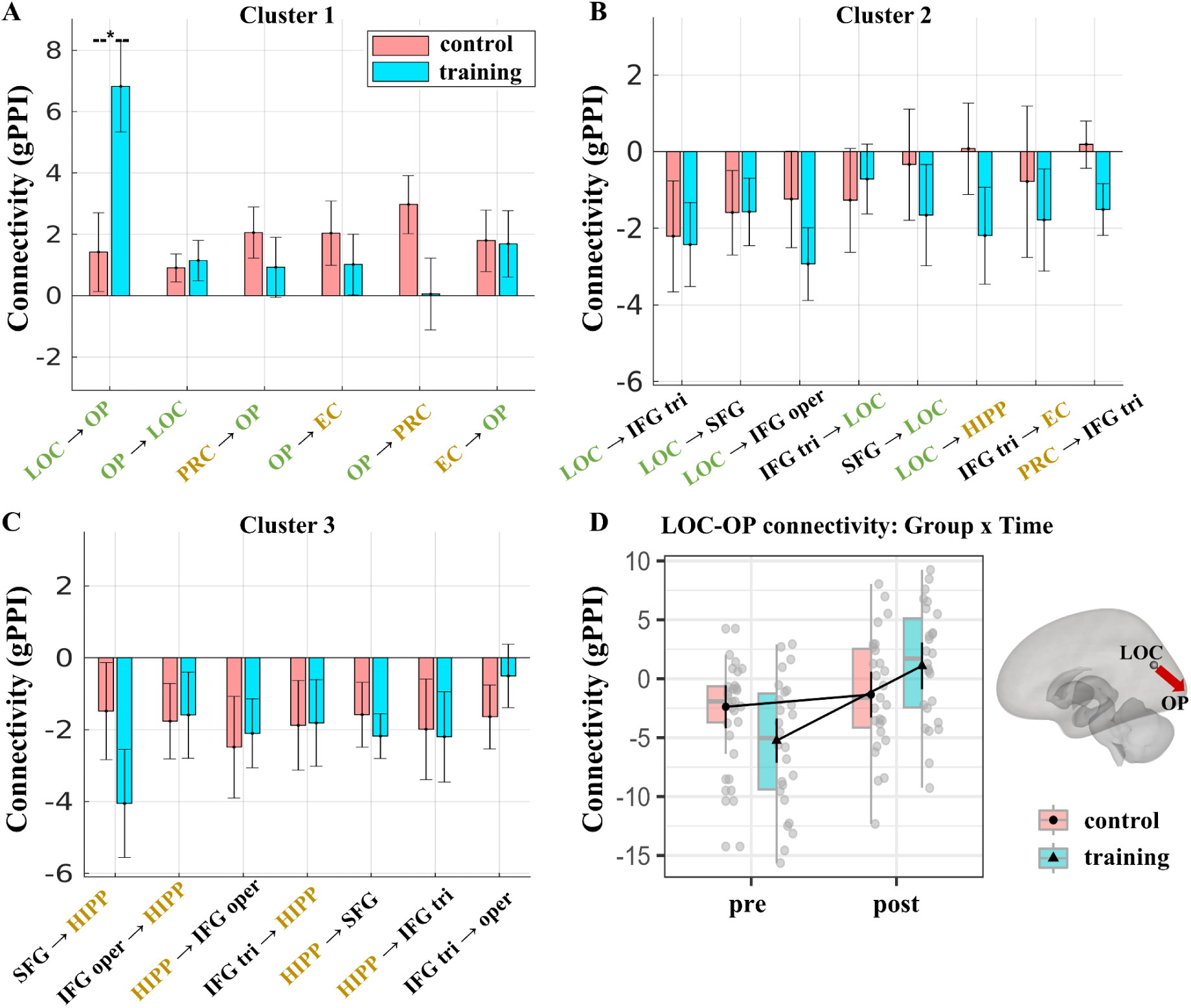
Brain connectivity training effects. **A-C**: Change in LD connectivity (post-training minus pre-training) in connections of clusters 1-3. **D**: Interaction plot of the Group x Time (pre-, post-training) effect (ANOVA) for the LOC-OP connection (lateral occipital cortex – occipital pole). We found a significant training-induced increase in task-based connectivity from the LOC to the OP. * p < .05 p-FDR corrected. **PRC**: perirhinal cortex, **EC**: entorhinal cortex, **HIPP**: hippocampus**, LOC**: lateral occipital cortex, **OP**: occipital pole**, SFG**: superior frontal gyrus, **IFG**: inferior frontal gyrus, **tri**: triangularis, **oper**: opercularis. The colour of the ROIs’ names denote whether they are a visual (green), MTL (gold), or PFC (black) area.

Focusing on the connections of cluster 1, the mixed ANOVAs demonstrated a significant Group x Time interaction effect in the LOC-OP connection (*F*(1,49) = 10.00, *p-FDR* = .016, η²_G_ = .092) (Table 6, Fig. 6D). Post-hoc analyses showed that the training group exhibited higher LOC-OP connectivity post- (M= 1.52, SE = 0.994) versus pre-training (M = −5.59, SE = 0.975) (*t*(49) = −5.12, *p* < .001), whereas the control group exhibited no significant difference post- (M = −1.40, SE = 0.951) versus pre-training (M = −2.44, SE = 0.932) (*t*(49) = 0.78, *p* = .439). Conversely, we found no significant training-induced effect in any other functional connection within the cluster 1, 2 or 3 (Tables S1-S3). Therefore, the cognitive-training intervention led to a selective effect of higher task-based LD functional connectivity from the LOC to the OP visual area, but no significant effects for hippocampal-PFC, visual-PFC or any visual to MTL task-based functional connections.

**Table 6.**
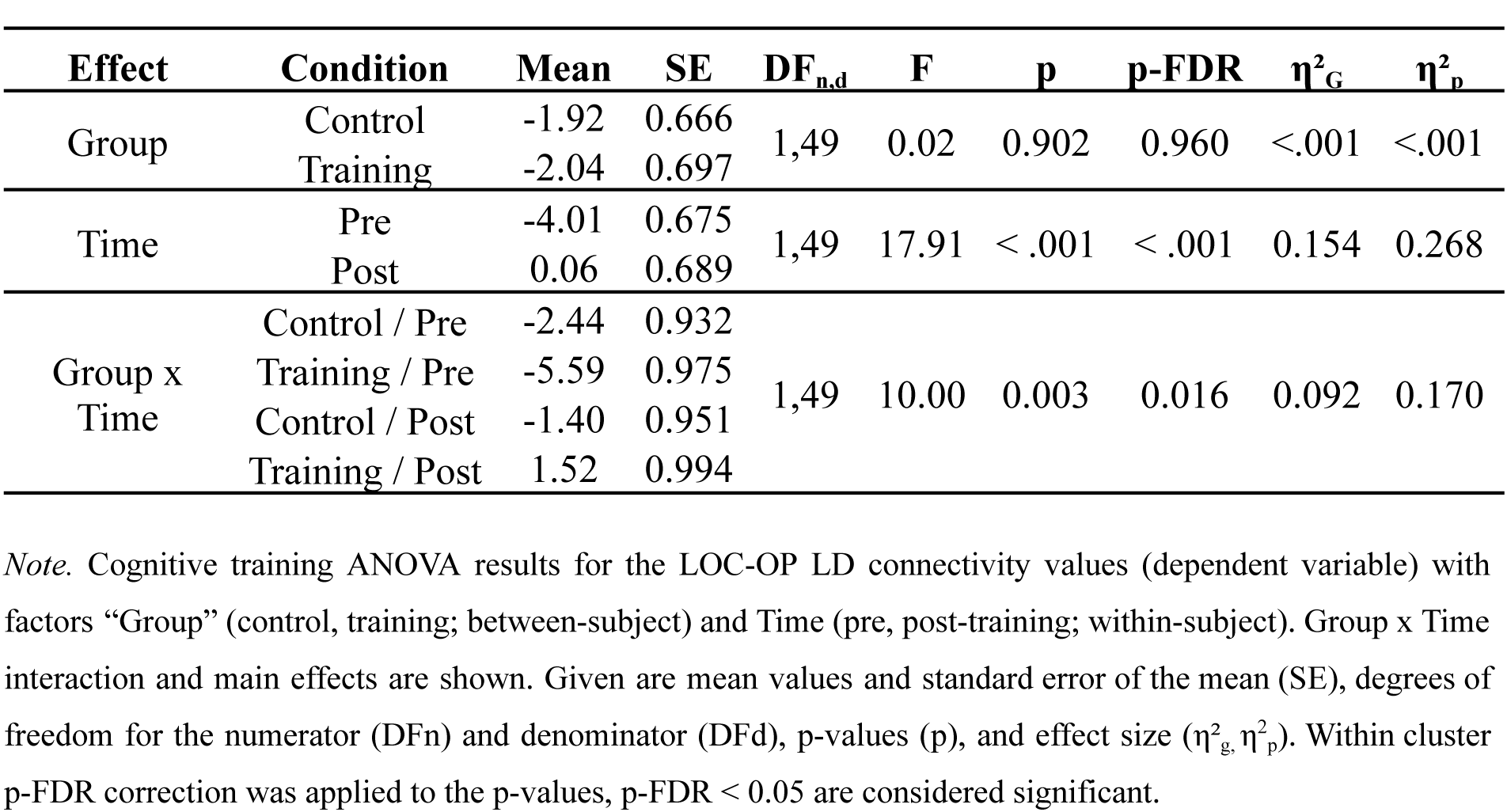
ANOVA cognitive training results for the LOC-OP LD connectivity.

### 2.4 Post-training hippocampal-PFC connectivity change is associated with behavioral improvement

Next, to test whether connectivity changes within the training group are linked to behavioral improvement, we fitted linear regression models to investigate the relationship between the post- versus pre-training change in LD connectivity and the A discriminability index (dependent variable) (covariates: age, sex). We focused on the connection LOC-OP (since it exhibited a group-level training effect) and the hippocampal-PFC composite connectivity score (since it was associated with inter-individual performance in the baseline data; see section 2.2).

The results revealed that decreases in LD hippocampal-PFC connectivity were strongly associated with greater behavioral improvement in the A index (p = .006, p-FDR = .018, R^2^_p_ = .291, b = −.031) (Fig. 7). To ensure this relationship is not disproportionately affected by outliers, we validated the above model using also robust regression (robust model: p = .002) (see Fig. S3). We observed no significant relationship between behavioral improvement and LOC – OP connectivity change (p = .782, R^2^_p_ = .003, b = .000), or between the hippocampal-PFC connectivity change and behavioral change in the lure discrimination index (LDI) at the MST task (p = .810, R^2^_p_ = .003, b = .012). Altogether, these results suggest that LD connectivity changes are linked to MD performance improvement, with decreased hippocampal-PFC connectivity being associated with greater improvement.

**Figure 7.**
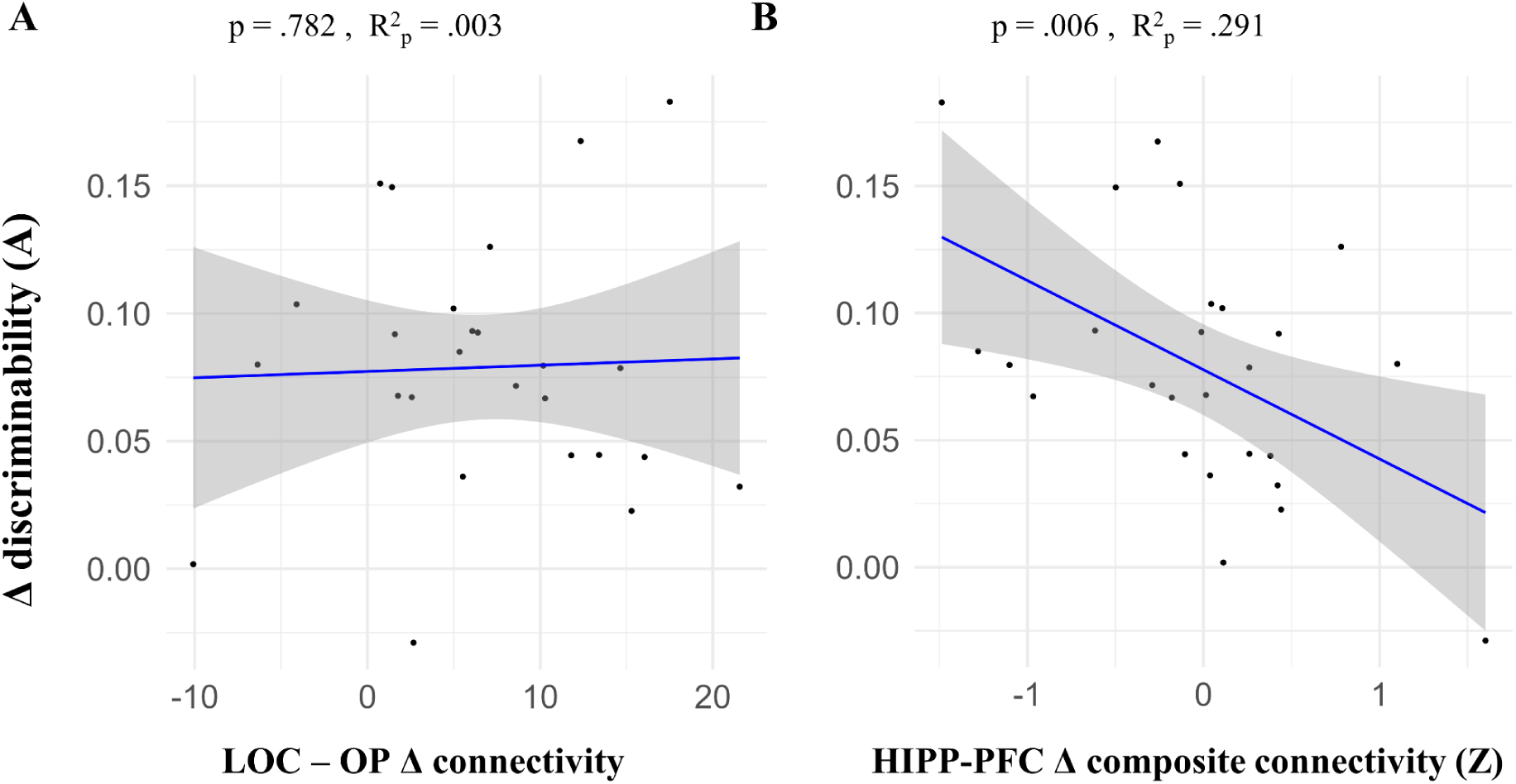
Brain connectivity change – behavior change (Δ: post-minus pre-training). Linear regression models fitted within the training group. X axis depicts the independent variable: LD connectivity Δ values (gPPI) of a connection (seed ROI – target ROI) or a composite score of a group of connections. Y axis shows the dependent variable (discriminability index A). P-values and partial R square (R^2^_p_) values are shown. R^2^_p_ is given to estimate the variance explained by each connection independently of the covariates. **LOC**: lateral occipital cortex, **OP**: occipital pole, **HIPP-PFC**: hippocampal-prefrontal connectivity composite score (Z score; see also section 2.2).

## 3. Discussion

We previously reported that MD training led to improved MD performance and regional functional changes (Güsten et al., 2024). Here we investigated how functional connectivity patterns of MTL, PFC and visual brain regions relate to mnemonic discrimination (MD) and interindividual differences in MD performance before and after cognitive training. We observed a lure detection (LD) functional connectivity signature involving MTL-PFC-visual areas (Fig. 3). Across individuals, higher LD task connectivity between the hippocampus and PFC areas was associated with poorer MD performance, whereby low performers showed higher LD connectivity (Fig. 4). After two weeks of MD training, stronger decreases in hippocampal-PFC LD connectivity were associated with greater MD improvement (Fig. 7). Finally, training led to greater LD connectivity from the lateral occipital cortex (LOC) to the occipital pole (OP) visual area (Fig. 6).

While the contributions of hippocampal and extra-hippocampal cortical regions to MD are recognized (Amer & Davachi, 2023), their functional interactions during MD remain poorly understood. Our study addressed this gap, demonstrating that task-based functional connectivity patterns within the MTL-PFC-visual network during LD are closely linked to successful MD (Fig. 3). Specifically, we found elevated LD connectivity between the hippocampus and PFC regions (including the inferior frontal gyrus pars triangularis, pars opercularis, and superior frontal gyrus) at the group level.

Our findings align with previous studies suggesting a role for PFC-hippocampus communication in MD (Johnson et al., 2021; Nash et al., 2021; Wais et al., 2017, 2018). Interestingly, however, higher LD-related connectivity in those hippocampal-PFC connections predicted poorer MD performance within our sample (Fig. 4). One explanation is that increased hippocampal–PFC connectivity may reflect neural inefficiency (Neubauer & Fink, 2009). Individuals with poorer MD performance might over-recruit PFC regions, in synchrony with the hippocampus, without experiencing a benefit for task performance. Our findings raise the possibility that over-connectivity with PFC regions could interfere with the hippocampus’s specialized role in fine-grained discrimination required for MD. This interference may be due to an introduction of neural noise (animal findings show that prefrontal projections modulate hippocampal signal-to-noise ratio (Malik et al., 2022)), a failure to suppress irrelevant information (M. C. Anderson & Green, 2001) or due to competition between PFC and hippocampus-based memory processing. Competition may be due to involvement of PFC working memory mechanisms that conflict with hippocampal MD processing (D’Esposito & Postle, 2015; Funahashi, 2017; Lara & Wallis, 2015).

Instances of over-recruitment without performance benefits have been reported in earlier studies (Poldrack, 2015). One framework explaining such a negative relationship is the balance between functional integration and segregation within neural networks. Excessive integration—that is, increased connectivity—can disrupt specialized processing, leading to decreased performance on tasks relying on specific neural circuits (Shine & Poldrack, 2018). In the case of MD, heightened hippocampal–PFC connectivity might reflect an imbalance where excessive integration interferes with hippocampal function. Thus, maintaining optimal segregation of the hippocampus may be key for successful MD, while over-integration with PFC networks could impair this process.

Importantly, this relationship between higher LD hippocampal-PFC connectivity and lower MD performance was observed both during the fMRI paradigm and in an independent MD task performed out-of-scanner: the mnemonic similarity task - MST (Stark et al., 2019) (Table 3-4). This suggests that hippocampal-PFC connectivity during MD-related processing may be a task-invariant individual difference marker indicative of MD performance. To our knowledge this is the first time that interindividual differences in brain function (task-functional connectivity) related to performance across two different MD tasks have been identified. Further causal research is warranted to elucidate the functional role of hippocampal-PFC connectivity in MD, potentially through manipulations that modulate prefrontal or hippocampal activity, such as transcranial magnetic stimulation or focussed ultrasound stimulation (Luber & Lisanby, 2014).

Given the limitations of fMRI in capturing the temporal neural dynamics of functional connectivity, more direct measures, such as electrophysiological recordings of neural oscillations, could provide valuable insights into the mechanisms underlying hippocampal-PFC communication in MD. Theta oscillations have been proposed as a potential mechanism orchestrating memory-related hippocampal-prefrontal interactions (K. L. Anderson et al., 2010; Gattas et al., 2023; Herweg et al., 2016; Siapas et al., 2005; Wang et al., 2021). Further electrophysiological research is needed to determine the extent to which hippocampal-PFC coupling via theta oscillations supports MD and to elucidate the directionality of this communication. Additionally, brain stimulation techniques targeting PFC theta oscillations could test the causal role of theta-mediated connectivity in MD and explore its potential to enhance cognitive performance by modulating hippocampal-PFC communication.

In addition to hippocampal-PFC interactions, our findings highlight the role of visual cortex connectivity during MD. Our findings suggest that successful MD relates to a cortical network encompassing both lower and higher-order visual areas, interconnected with prefrontal and extra-hippocampal MTL regions (Fig. 3, Table 1). We interpret that these connectivity patterns support perceptually fine-grained, high-fidelity representations necessary for discriminating similar memories (Wais et al., 2017). Our findings extend previous studies showing the involvement of occipital visual areas in MD (Klippenstein et al., 2020; Pidgeon & Morcom, 2016) with new data on their functional connectivity. This inter-cortical communication may facilitate cortical pattern separation by fine-tuning visual input to the hippocampus and regulating the influence of top-down prefrontal control (Amer & Davachi, 2023; Gilbert & Li, 2013; Pidgeon & Morcom, 2016). This interpretation is consistent with research highlighting the role of top-down influences on visual processing, including the modulation of early visual areas by higher-order regions (Gilbert & Li, 2013). Our findings further highlight the relevance of extra-hippocampal contributions to MD, particularly within the visual processing hierarchy.

Our 2-week cognitive training intervention had an impact on early visual connectivity. We observed an increased connectivity from the LOC to the OP after training, suggesting enhanced early visual top-down feedback (Fig. 6, Table 6). This suggests that MD improvements may be partly driven by more fine-tuned input to the hippocampus and thus more effective visual cortical contributions to pattern separation (Amer & Davachi, 2023; A. M. C. Kelly & Garavan, 2005; Pidgeon & Morcom, 2016). This interpretation aligns with previous research demonstrating plasticity of visual cortical areas and presumably more refined representations following training (Green & Bavelier, 2008; Schwartz et al., 2002), including functional reorganization in the early visual cortex (Schwartz et al., 2002). The LOC is implicated in higher-order visual processing and object recognition (Grill-Spector et al., 2001) while the OP in early visual processing (Mulckhuyse et al., 2011). Enhanced visual representations, with sharper tuning to relevant features, could facilitate downstream mnemonic processes by providing the hippocampus with more distinct inputs. We note, however, that the physiological role of training-related LOC-OP visual connectivity changes for MD remains uncertain given the absence of a significant change-change correlation (Fig. 7).

Although we did not observe a significant training-related change at the group level, individuals who showed a post-training decrease in hippocampal-PFC connectivity also exhibited greater MD improvements (Fig. 7). Thus the same connectivity pattern that explained baseline performance variability, also explained inter-individual differences in performance improvements after training. This reinforces the idea that hippocampal-PFC connectivity is a neural resource for MD performance that not only explains individual differences but can also be targeted to improve performance. Training may have promoted a more efficient division of labor between PFC and hippocampus, where the hippocampus performed pattern separation more effectively with less reliance on top-down control from the PFC. Studies have shown that training can decrease PFC engagement, indicating increased processing efficiency (Vermeij et al., 2017). These change-change correlations were specific to our MD task because there was no correlation with MST performance change after training (also see (Güsten et al., 2024)). Compatible with this absence of transfer, we found no correlation between hippocampal-PFC connectivity and MST performance change. Given that at baseline the hippocampal-PFC LD connectivity was correlated with performance in both tasks, our MD task and the MST, we would have expected that change-change correlations after training would also be present for both tasks. One reason for this negative finding may have been the short training duration of two weeks.

While our 2-week training protocol revealed significant changes in visual cortex connectivity, longer interventions may be necessary to induce more widespread network changes. Future research should investigate whether extended training leads to broader functional reorganization, including changes in hippocampal-PFC connectivity, and whether these changes are associated with MD improvement transfer to non-trained MD tasks and other episodic memory tasks. Our study focused on young adults and so paves the way for future studies to investigate the efficacy and neural correlates of similar training in older adults, particularly those experiencing cognitive decline. Understanding how these training programs affect brain function in the context of aging and neurodegeneration is crucial for developing effective interventions to maintain cognitive health in older adults.

By using a memory training intervention, our study provided some causal evidence for the role of brain connectivity in MD. To further strengthen causality, future research utilizing brain stimulation techniques can be helpful. For instance, applying inhibitory stimulation to the PFC during MD tasks could help determine whether disrupting PFC activity leads to improved performance in individuals who initially exhibit high hippocampal-PFC connectivity.

In conclusion, we have identified a brain connectivity signature of MD that generalizes across two independent tasks. This neural signature also explained individual performance gains after training. Thus, we identified a neural resource for MD performance that not only explains individual differences but can also be targeted to improve memory performance.

## 4. Materials and Methods

### 4.1. Participants

Sixty young adults were recruited at the Otto von Guericke University of Magdeburg. Five participants voluntarily chose to discontinue with the study and one was excluded due to problematic brain extraction (M age = 23.76 years, SD = 3.35, 61.11% female). One further subject was excluded from the connectivity training analysis due to incomplete fMRI data, resulting in a total sample of 54 for our baseline analyses and 53 for our 2×2 training analyses. All participants were screened for prior psychiatric or neurological diagnoses and had intact or corrected vision. All participants provided written informed consent and were compensated for their time. The study was approved by the Otto von Guericke University Magdeburg ethics committee and conducted in accordance with the declaration of Helsinki, 2013.

#### 4.2.1 Study design, procedure and experimental paradigm

We used a 2×2 longitudinal intervention design (Group x Time). Participants were randomly assigned to an experimental training group (n=26) or an active control group (n=27) with a task fMRI session completed pre- and post-a 2-week computerized web-based cognitive training intervention (Fig. 1, 8). Pre- and post-training, participants performed at the scanner, while fMRI data was collected, a 6-back version of the object-scene task developed to measure MD (Berron et al., 2018; Güsten et al., 2021) (see Fig. 1D). Before scanning, subjects were given standardized instructions and underwent a short training session. Stimuli were presented via a mirror on a magnetic resonance (MR) compatible display. When necessary, vision was corrected using MR-compatible glasses.

The object-scene task is a continuous old/new recognition task (Fig. 1A-C), described in detail in Berron et al. (2018). In this task, subjects have to recognize whether each item presented is a ‘new’ or an ‘old’ (repeated) image by responding with their right index or middle finger. We adapted it for our study by using a 6-back instead of the original 2-back format, to boost difficulty given participants’ young age and expected post-training task proficiency. The task had 2 x 13 mins runs. Stimuli were presented in sequences of 12 items each (6-back format: 6 items presented in encoding, 6 in test phase). A sequence consisted of either object or scene stimuli only presented on a gray background. In each sequence, the first 6 stimuli were shown for the first time (‘first’ trials; correct response: ‘new’), while each of the following six stimuli (test phase) could be either an exact repetition (‘repeat’ trial; correct response: ‘old’) or a very similar yet different version of the first images (‘lure’ trial; correct response: ‘new’). In lure trials, local features of the objects or global features (e.g., the geometry) of the scenes were changed (Fig. 1C), but not the color, position, size or viewpoint. Ten gray noise images (‘scrambled’) were presented as perceptual control trials at the start and end of each run (40 trials in total). In total, 60 sequences were shown. This resulted in a 2×2 factorial design with 30 trials for each trial type combination (lure objects, repeats object, lure scenes, repeats scenes). The sequences’ order was counterbalanced with respect to the object - scene, and repeat - lure trial types. The stimuli were presented in an event-related design with each stimulus shown for 3 seconds (secs), being separated by a fixation star. Jittered inter-stimulus intervals were used (range: 0.8 to 4.2 secs, mean ∼ 1.63 secs) to optimize statistical efficiency (Dale, 1999). The task difficulty (correct response rate) for each image was validated in an independent study (Güsten et al., 2021), ensuring that different stimuli sets of similar difficulty were used at the two fMRI sessions. Note that the stimuli used for the cognitive training were created using the same principles.

#### 4.2.2 Cognitive training

In between fMRI scans, all participants performed a 2-week computerized cognitive training on an online platform (http://iknd-games.ovgu.de/MemTrain/), comprising 6 training sessions in total (Fig. 8). The training group performed the 6-back object-scene task described above. The stimuli were chosen from items assessed in Güsten et al. (2021). The training consisted of 45 different levels in total. Subjects progressed to the next level upon reaching a threshold of correct responses (ranging from 70 % up to 90 % across the levels), otherwise they would repeat a level until they achieved the threshold. At each level, the set of stimulus pairs was fixed, and the presentation of each stimulus version (1 or 2) was randomized for both the presentation and test trial. The levels were created with increasing difficulty as one progressed in the training. The task consisted of rounds. Each round was made up of 4 blocks (2 object and 2 scene blocks). Each block contained a presentation and test phase. First, 6 stimuli were presented for 3 secs each, separated by a blank page (1.5 secs). Then, a countdown (3 secs) indicated the test trials, in which participants had to indicate whether a presented stimulus was a repeat or a lure (max. 4.5 secs). A training session ended after spending 45 minutes in the training rounds.

**Figure 8.**
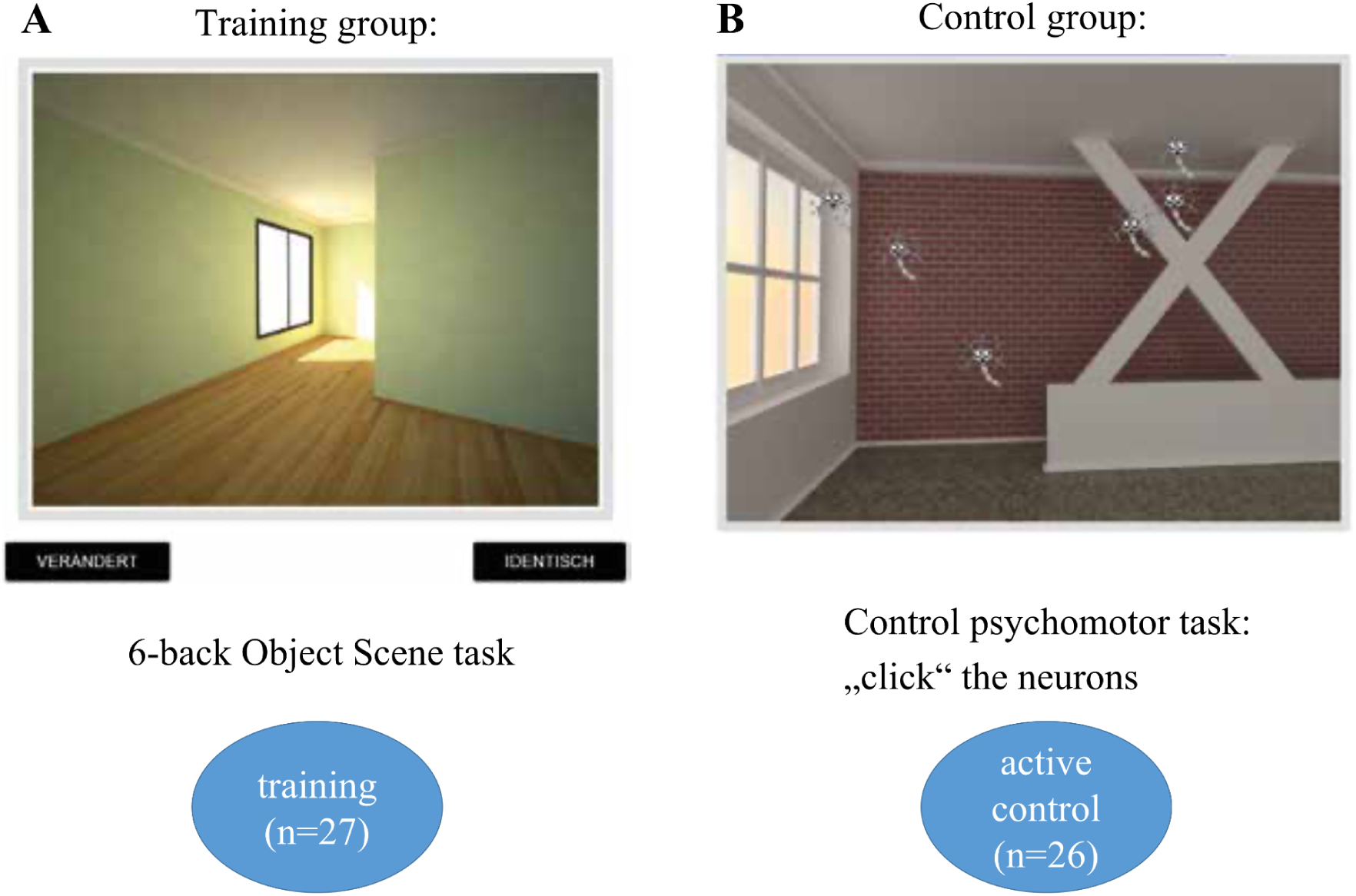
Training design overview. Participants underwent a computerized training, from a remote location (mobile). Both training and control groups underwent active training, but on different tasks. **A.** The training group performed a 6-back version of the object-scene task (Berron et al., 2018; Güsten et al., 2021), with separate object and scene trials, and domain-specific level increase. Level difficulty was increased by showing more difficult images based on previously evaluated difficulty. Stimulus discriminability rather than trial length was modified to more directly target discrimination ability**. B.** The control group training consisted of eliminating moving neurons by clicking on them. With level increase, neurons became more numerous and faster. To keep stimuli visually similar to the main task, background images were equivalent to the scenes from the training task. Figure modified from Güsten et al (2024). For more details on the training see Güsten et al. (2024).

The control group trained on a psychomotor task in which they had to click on small moving neuron-icons presented on top of background images (see Fig. 8). To keep the visual stimulation similar to the actual training task, the background images were pseudorandomly taken from the stimulus set of the training group. The training structure was the same as in the main training task to keep the training flow comparable. Namely, one training round consisted of 4 blocks: 2 blocks with objects and 2 blocks with scene stimuli appearing in the background. A higher level was reached by eliminating a certain percentage (varied from 80 to 90 % across the levels) of neurons within a round. Again, a training session ended after spending 45 minutes in the training rounds. The control task was progressively made more challenging at higher levels in several ways: 1) reducing the neurons’ presentation time, 2) increasing the number of neurons to eliminate, 3) increasing neurons’ moving speed, 4) making neurons change direction and 5) introducing ‘bad’ neurons, which participants had to avoid clicking.

#### 4.2.3 Out-of-the scanner cognitive tasks

In addition to the experimental paradigm performed in the scanner, all subjects completed a series of transfer tasks as explained in detail in (Güsten et al., 2024). Here we examined whether task-related connectivity in our MD task, which was performed at the scanner, is associated with a series of independent memory tasks performed outside of the scanner at another day (part of the transfer tasks) (see Table 4). Those memory tasks included the mnemonic similarity task (MST) (Stark et al., 2013, 2019), a modified 30-word list verbal learning memory task (Helmstaedter et al., 2001), the Rey-Osterrieth Complex Figure (ROCF) (Osterrieth, 1944), and the object-in-room recall (ORR) task (Berron et al., 2024).

The MST (Version 0.9) (Stark et al., 2013) assessed mnemonic discrimination through separate encoding and retrieval phases. In the encoding phase, participants made indoor-outdoor judgments for 128 object stimuli. The retrieval phase involved identifying 192 stimuli (64 repeats, 64 similar lures, and 64 foils) as either repeats, lures, or foils. The LDI and corrected hit rate measures were calculated. The LDI measures the ability to detect lures while controlling for response bias. The corrected hit rate estimate (“old”|old - “old”|lure) is an alternative bias-corrected measure of discrimination (Loiotile & Courtney, 2015). To assess memory retrieval and pattern completion, the ORR task was employed. The ORR task (a digital version of the task is reported in (Berron et al., 2024)) involved participants memorizing rooms containing objects and their specific locations. It consisted of an encoding phase, where participants memorized 75 room-object combinations, followed by an immediate retrieval test to ensure accurate encoding. During retrieval, participants viewed a room with a cued location and selected the correct object from three options: the correct object, an object previously present in the picture but in a different location (internal lure) and an object not present in the room at all (external lure).

To assess visual memory, visuospatial constructional ability, and recognition, the Rey-Osterrieth Complex Figure (ROCF) was used. Participants first copied the figure, then reproduced it from memory after 3 minutes, and again approximately 30 minutes later. For verbal long-term memory and interference, a modified version of the Verbal Learning and Memory Test (VLMT) was applied, the lists were extended to a length of 30 words by adding the lists from the CVLT. Participants were exposed to lists A and C, with interference introduced through the VLMT’s 15-word list, followed by list D in the post-test. The procedure involved the experimenter reading the word list five times, each followed by immediate recall (L 1-5), then an interference list recall, a recall of the original list (L 6), and a final recall (L 7) after a 30-minute delay. For a more detailed description of the task and associated measures, see also (Güsten et al., 2024).

### 4.3 Imaging data acquisition

All imaging data were acquired on a 3T MAGNETOM Skyra scanner (Siemens, Erlangen, Germany) using syngo MR E11 software and a 64-channel head coil. Structural images were acquired using a T1-weighted MPRAGE sequence with 1 mm isotropic resolution (3D acquisition; TR/TE/TI = 2500/4.37/1100ms, FoV=256 x 256mm²; flip angle= 7 degrees; BW = 140Hz/Px). Whole-brain functional data were acquired in 2 runs of 13 minutes each, using T2*-weighted echo planar imaging (EPI). We used a 2D simultaneous multi-slice EPI sequence (SMS-EPI) developed at the Center for Magnetic Resonance Research, University of Minnesota (CMRR; Moeller et al., 2010), with a 2 x 2mm² resolution, FOV = 212 x 212mm², TR/TE = 2200/30ms, 10% slice gap, multiband acceleration factor 2, GRAPPA 2, phase encoding (PE) direction P > A, 64 slices and 2mm slice thickness. In addition, phase maps were acquired with the following parameters: TR/TE1/TE2 = 675/4.92/7.38 ms, spatial resolution = 3 mm, and 48 slices.

### 4.4. MRI preprocessing and denoising

We processed MRI data using the ‘fMRIPrep’ (v. 20.2.6) pipeline (Esteban et al., 2019), employing default options, and the CONN toolbox (Whitfield-Gabrieli & Nieto-Castanon, 2012). Each T1-w image was corrected for intensity non-uniformity and skull-stripped. A T1w-reference map was computed by registering 2 T1w images acquired from each subject (which were collected within a 2-week period) to obtain a robust estimate of anatomy. Spatial normalization to the MNI space (ICBM 152 Nonlinear Asymmetrical template 2009c) was performed through nonlinear registration. Tissue segmentation was performed on the brain-extracted T1w images. The functional data were slice-time corrected and co-registered to the T1w using boundary-based registration with 9 degrees of freedom. Physiological noise regressors were extracted using CompCor, in addition to other confound variables, including head-motion parameters and framewise displacement (FD), which were used later at denoising. Frames that exceeded a threshold 0.5 mm FD were annotated as motion outliers. All resampling was performed in a single interpolation step. For further details including the software used in each step, refer to the documentation: https://fmriprep.readthedocs.io/en/20.2.6/.

Next, the functional data were smoothed using a 4 mm FWHM Gaussian kernel (determined a-priori as twice the voxel size in order to compromise anatomical specificity and signal-to-noise ratio increase), and denoised using the CONN toolbox to remove potential confounding effects in the BOLD signal. The denoising steps included: regressing out 3 translation and 3 rotation motion parameters plus their first-order derivatives (12 parameters in total), anatomical component-based noise correction procedure (aCompCor) to remove noise components from the white matter and cerebrospinal areas, scrubbing to remove the volumes identified as outliers due to excessive motion (> 0.5 mm FD), high-pass filtering 1/128 Hz, linear detrending, and despiking. Covariates were also included to account for potential slow trends, initial magnetization transients, or constant task-induced responses in the BOLD signal, according to the default denoising pipeline in CONN (Nieto-Castanon, 2020). For further details, see the documentation: https://web.conn-toolbox.org/fmri-methods/denoising-pipeline.

### 4.5 Functional connectivity analyses

#### 4.5.1. Regions of interest (ROIs)

We performed hypothesis-driven region of interest (ROI)-to-ROI analysis. For each ROI, unsmoothed BOLD data was extracted by default in CONN. We constructed a model including major medial temporal lobe (MTL), prefrontal (PFC) and visual ROIs: hippocampus, enthorhinal cortex (EC), perirhinal cortex (PRC), parahippocampal cortex (PaHC), superior frontal gyrus (SFG), middle frontal gyrus (MidFG), and inferior frontal gyrus (IFG) including pars triangularis (tri) and pars opercularis (par), medial prefrontal cortex (medFC), lateral occipital cortex (LOC) and occipital pole (OP) (Fig. 2). All ROIs were bilateral at MNI152 space. The EC was extracted from the Juglich atlas (Eickhoff et al., 2005), the PRC and PaHC from established MNI-space segmentations (Ritchey et al., 2015) (https://neurovault.org/collections/3731/). The remaining ROIs were derived from the Harvard-Oxford cortical and subcortical atlases (Desikan et al., 2006).

#### 4.5.2 Task-modulated connectivity metric (gPPI) and MD contrast

We applied a region of interest generalized psychophysiological interaction (gPPI) analysis (McLaren et al., 2012) to examine the task-modulated functional connectivity between the ROIs in our model (ROI-to-ROI analyses) for the successful MD contrast (*lure detection - LD*: correct lure minus repeat trials). All functional connectivity analyses performed in our study refer to this LD contrast. We used the CONN toolbox (Nieto-Castanon, 2020) for all analyses. gPPI values were calculated for each ROI-to-ROI connection pair for each participant, representing LD task functional connectivity from a seed to a target region. gPPI measures the influence of a seed on a target region after partialling out task-related activity and task-unrelated connectivity, resulting in an asymmetrical effective connectivity matrix. This means that each connection given follows a seed – target format that denotes directionality.

#### 4.5.3 Characterisation of baseline LD profile

To typify the connectivity patterns associated with successful MD, we conducted analyses of the LD contrast in the pre-training whole-sample data, without respect to subsequent group allocation. This resulted in an 11 x 11 gPPI matrix (11 ROIs, 110 connections). We used a cluster-level inference with the default ROI-to-ROI network multivariate parametric statistics approach in CONN.. This approach uses multivariate statistics to examine groups of related connections, resulting in an F-statistic for each cluster with FDR-corrected cluster-level p-values, in addition to a post-hoc connection-level thresholding that keeps the strongest connections within each significant cluster. We used the standard settings: p < .05 cluster-level p-FDR, connection threshold: p < .05 p-uncorrected.

#### 4.5.4 Cognitive-training effects on LD connectivity

To test the training effects on the ROI connectivity data, we performed mixed 2 x 2 analysis of variance (ANOVA) (aov_ez function default type III sum of squares option, R package ‘afex’ v. 1.3-0) (Singmann et al., 2023). As the dependent variable, we used the gPPI connectivity values from each connection in each of the clusters identified from the previous baseline characterization step. We added the Group (training, control: between-subjects factor) x Time (pre-, post-training: within-subjects factor) as independent variables, the sex and age as covariates. We focused on the Group x Time interaction effect to probe the training effects. We applied Hyunh-Feldt correction for sphericity violations cases, and calculated both the generalized Eta-squared (η²_G_) and partial Eta-squared (η²_p)_ to estimate the effect size for the ANOVAs (Lakens, 2013). Post-hoc tests were conducted with the emmeans R package v. 1.8.8 (Lenth, 2023). When required, simple effects and an interaction contrast were calculated to test if the training-induced (post- versus pre-training) change was bigger for the training compared to the control group. We applied p-FDR multiple comparisons correction for the tests performed within each cluster of connections to strike a statistical power and control balance.

### 4.6 Behavioral data processing and cognitive-training effects

The behavioral data from the MD fMRI task were processed using R (v. 4.3.1) (R Core Team, 2023). We calculated 4 readouts; hit rates (repeats correct), correct rejection rates (lures correct), false alarm rates (lures incorrect), and the *A* discriminability index (a non-parametric discriminability estimate based on the hits minus false alarms rate and uses a corrected formula to estimate the non-parametric sensitivity associated with the signal and noise distributions) (Zhang & Mueller, 2005). This *A* index is regarded as an improved alternative to A prime. We calculated total scores by combining both modalities (objects, scenes), as we were interested in the total modality-independent MD performance.

To test the training effect on MD performance we employed 2×2 mixed ANOVAs (Factors: Group x Time, covariates: sex, age) following the same statistical approach as for the ROI connectivity data (see section 4.5.4). We used the A discriminability Index as the main dependent variable, but also examined the underlying trial types separately: correct rejections (correct lures) and hits (correct repeats).

#### 4.7.1 LD connectivity – behavior regression

To examine the LD connectivity-behavior relationship, we fitted linear regression models in the pre-training data (‘lm’ R function). Each model used the formula: A ∼ connectivity + age + sex (A discriminability: dependent variable. Connectivity: values of a certain connection as the independent variable). Age and sex were included as covariates of no interest. We applied p-FDR multiple comparisons correction for the tests performed within each cluster of connections. To estimate the variance explained by a connection we calculated the partial R square (R^2^_p_) using the ‘sensemark’ R package v. 0.1.4 (Cinelli et al., 2020).

#### 4.7.2 Relationship between LD hippocampal-PFC connectivity and other memory tasks

To test the association between LD task connectivity in the hippocampal-PFC network and performance on various out-of-the scanner independent memory tasks, we conducted a multivariate regression analysis using the LD hippocampal-PFC composite connectivity score as the main predictor. This score was calculated as the average of the z-score transformed connectivity values extracted from each of the six hippocampal-PFC connections values (which belong to the cluster 3 and were also found to be linked to high performance). Dependent variables included performance scores from multiple memory tasks including the MST task (measures: LDI and corrected hit rate), the ORR task (measures: correct retrieval performance, and retrieval internal errors), the Rey-Osterrieth Complex Figure (measure: delayed recall), and VLMT test (word list test: encoding vs delay recall score (List5 minus List7 performance)). Age and sex were included as covariates. The Pillai’s statistic was calculated to assess the overall model fit and determine the significance of the connectivity score effect across the different memory measures using the ‘Anova’ function (‘car’ package) and the ‘lm’ function in R. Then we examined using linear models the relationship of the hippocampal-PFC composite connectivity score with each separate memory measure (dependent variable).

#### 4.7.3 LD connectivity change – behavior change regression

To examine the LD connectivity-behavior change (Δ) relationship (post- minus pre-training), we fitted linear regression models (‘lm’ R function) within the training group; we focused on the LOC-OP connectivity (for which we observed a group training effect), and hippocampal-PFC composite connectivity score (which was associated to MD performance at the baseline data). Each model used the ‘A Discriminability Δ’ as the dependent variable, the ‘connectivity Δ’ as the independent variable, while ‘age’ and ‘sex’ were included as covariates of no interest. To estimate the variance explained by the connectivity change, we calculated the partial R square. To ensure that outliers did not disproportionately affect the hippocampal-HPC connectivity-behavior change results, we further validated our model using robust linear regression that is less sensitive to outliers (Fig. S3) (MM-type estimator; “lmrob” function from the “robustbase” R package, version 0.99-4-1) (Maechler et al., 2024).

## Acknowledgments

This work was supported by Deutsche Forschungsgemeinschaft (DFG, German Research Foundation) – Project-ID 425899996 – SFB 1436. We are grateful to Aditya Nemali for his help in setting up the training website. We thank all participants who volunteered to participate in the training. We also want to thank the staff at the IKND and the university clinic for neurology, Medical Faculty, Otto-von-Guericke University for assistance in testing and scanning the participants.

## Abbreviations

AD: Alzheimer’s disorder
CL: correct lures
Δ: Delta (change post versus pre-training)
fMRI: functional magnetic resonance imaging
gPPI: generalized psychophysiological interaction
HIPP: hippocampus
IFG: inferior frontal gyrus;
tri: triangularis
oper: opercularis
LD: lure detection
LDI: lure discrimination index
LOC: lateral occipital cortex
MD: mnemonic discrimination
MedFC: medial frontal cortex
MidFG: middle frontal gyrus
MTL: medial temporal lobe
MST: mnemonic similarity task
OP: occipital pole
ORR: object-in-room recall
PaHC: parahippocampal cortex
PFC: prefrontal cortex
PRC: perirhinal cortex
Reps: repeats
ROI: region of interest
ROCF: Rey-Osterrieth Complex Figure
SFG: superior frontal gyrus
VLMT: verbal learning and memory test

## Supplementary information

### Results

#### S2.1 Behavioral data at baseline

**Figure S1.**
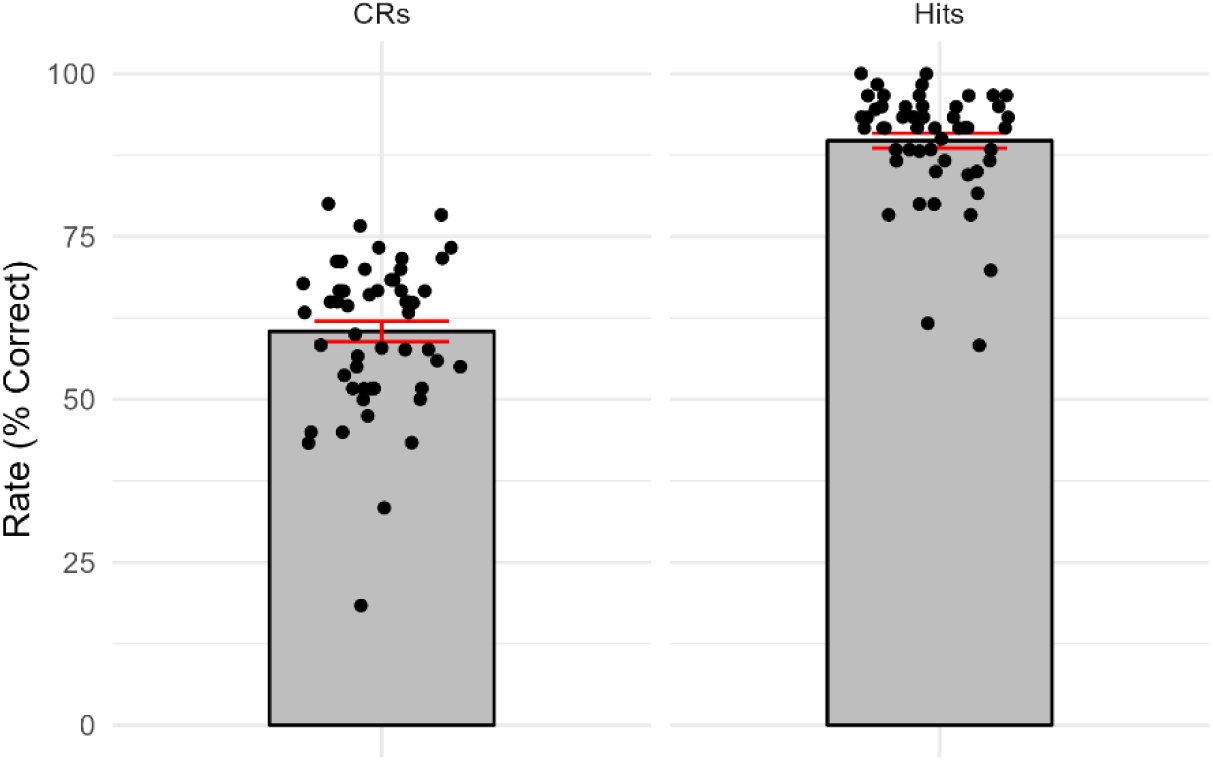
Behavioral performance at baseline in the object-scene MD task. The performance (percent correct) is plotted separately for the correct repeats (hits) and the correct lure trials (CRs – correct rejections) for the whole sample (*n = 54*). All data points correspond to the baseline (pre-training) time point.

**Figure S2.**
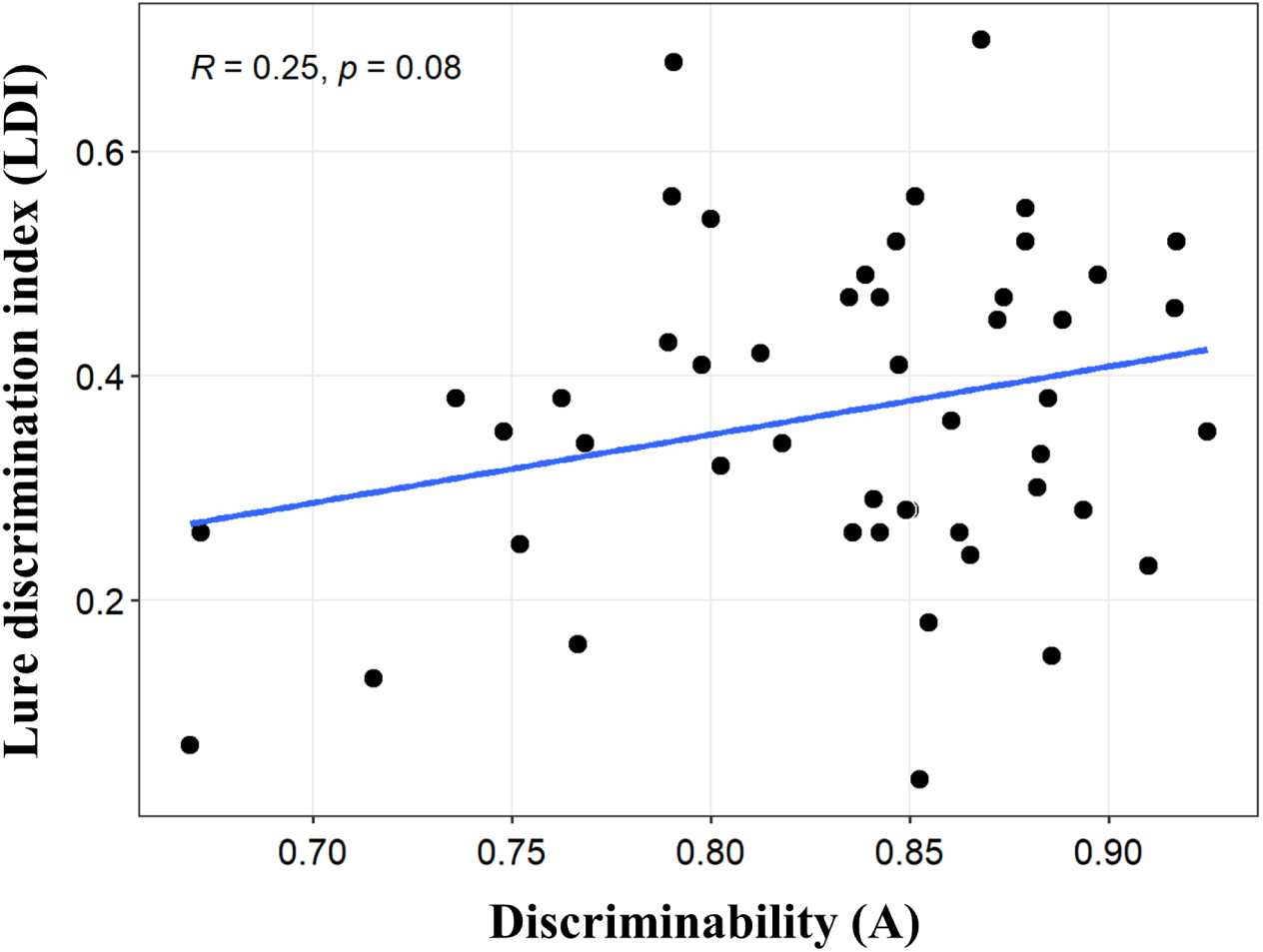
Correlation between baseline performance on the two different mnemonic discrimination tasks. The X axis represents performance on the object-scene MD task (discriminability A), while the Y axis represents performance on the MST task (LDI index) for the whole sample (*n = 54*). Both measures are bias-corrected and derived from their respective MD tasks. All data points correspond to the baseline (pre-training) time point*. R =* correlation index (Pearson r), *p* = p-value.

#### S2.2 ANOVA cognitive training connectivity

**Table S1.** ANOVA cognitive training results for connectivity values in cluster 1.

| Connection | Condition | Mean | SE | DF<sub>nd</sub> | F | p | p-FDR | $\eta^2_G$ | $\eta^2_p$ |
| --- | --- | --- | --- | --- | --- | --- | --- | --- | --- |
| LOC-OP | Control / Pre | -2.44 | 0.93 | 1,49 | 10.00 | **0.003** | **0.016** | 0.092 | 0.170 |
|  | Training / Pre | -5.59 | 0.98 |  |  |  |  |  |  |
|  | Control / Post | -1.40 | 0.95 |  |  |  |  |  |  |
|  | Training / Post | 1.52 | 0.99 |  |  |  |  |  |  |
| OP-LOC | Control / Pre | -1.40 | 0.39 | 1,49 | 0.19 | 0.663 | 0.796 | 0.002 | 0.004 |
|  | Training / Pre | -1.12 | 0.41 |  |  |  |  |  |  |
|  | Control / Post | -0.57 | 0.44 |  |  |  |  |  |  |
|  | Training / Post | 0.07 | 0.46 |  |  |  |  |  |  |
| PRC-OP | Control / Pre | -1.67 | 0.75 | 1,49 | 0.41 | 0.525 | 0.787 | 0.003 | 0.008 |
|  | Training / Pre | -1.31 | 0.78 |  |  |  |  |  |  |
|  | Control / Post | 0.29 | 0.68 |  |  |  |  |  |  |
|  | Training / Post | -0.21 | 0.71 |  |  |  |  |  |  |
| OP-EC | Control / Pre | -1.69 | 0.66 | 1,49 | 0.44 | 0.509 | 0.787 | 0.004 | 0.009 |
|  | Training / Pre | -1.05 | 0.69 |  |  |  |  |  |  |
|  | Control / Post | 0.54 | 0.78 |  |  |  |  |  |  |
|  | Training / Post | 0.20 | 0.81 |  |  |  |  |  |  |
| OP-PRC | Control / Pre | -2.30 | 0.88 | 1,49 | 2.74 | 0.104 | 0.313 | 0.021 | 0.053 |
|  | Training / Pre | -0.33 | 0.92 |  |  |  |  |  |  |
|  | Control / Post | 0.61 | 0.85 |  |  |  |  |  |  |
|  | Training / Post | 0.03 | 0.89 |  |  |  |  |  |  |
| EC-OP | Control / Pre | -1.42 | 0.83 | 1,49 | 0.01 | 0.927 | 0.927 | <.001 | <.001 |
|  | Training / Pre | -1.53 | 0.87 |  |  |  |  |  |  |
|  | Control / Post | 0.48 | 0.75 |  |  |  |  |  |  |
|  | Training / Post | 0.50 | 0.79 |  |  |  |  |  |  |

*Note.* ANOVA results examining the effect of cognitive training (Group x Time interaction) on connectivity for each individual connection (factors ‘Group’: control, training; between-subject. ‘Time’: pre-, post-training; within-subject). Given are mean values and standard error of the mean (SE), degrees of freedom for the numerator (DFn) and denominator (DFd), p-values (p), and effect size ( $\eta^2_G$ , $\eta^2_p$ ). Within cluster p-FDR correction was applied to the p-values. P-FDR < 0.05 are considered as significant.

**Table S2.** ANOVA cognitive training results for connectivity values in cluster 2.

| Connection | Condition | Mean | SE | DF<sub>n,d</sub> | F | p | p-FDR | $\eta^2_G$ | $\eta^2_p$ |
| --- | --- | --- | --- | --- | --- | --- | --- | --- | --- |
| LOC-IFG tri | Control / Pre | 2.01 | 0.794 | 1,49 | 0.14 | 0.710 | 0.911 | 0.002 | 0.003 |
|  | Training / Pre | 2.66 | 0.831 |  |  |  |  |  |  |
|  | Control / Post | 0.23 | 0.744 |  |  |  |  |  |  |
|  | Training / Post | 0.20 | 0.779 |  |  |  |  |  |  |
| LOC-SFG | Control / Pre | 1.61 | 0.746 | 1,49 | 0.01 | 0.911 | 0.911 | <.001 | <.001 |
|  | Training / Pre | 1.60 | 0.780 |  |  |  |  |  |  |
|  | Control / Post | 0.22 | 0.559 |  |  |  |  |  |  |
|  | Training / Post | 0.05 | 0.584 |  |  |  |  |  |  |
| LOC-IFG oper | Control / Pre | 0.56 | 0.758 | 1,49 | 1.69 | 0.200 | 0.534 | 0.020 | 0.033 |
|  | Training / Pre | 2.49 | 0.793 |  |  |  |  |  |  |
|  | Control / Post | -0.43 | 0.678 |  |  |  |  |  |  |
|  | Training / Post | -0.57 | 0.709 |  |  |  |  |  |  |
| IFG tri-LOC | Control / Pre | 1.44 | 0.711 | 1,49 | 0.02 | 0.903 | 0.911 | <.001 | <.001 |
|  | Training / Pre | 1.77 | 0.744 |  |  |  |  |  |  |
|  | Control / Post | 0.55 | 0.723 |  |  |  |  |  |  |
|  | Training / Post | 1.08 | 0.757 |  |  |  |  |  |  |
| SFG-LOC | Control / Pre | 1.26 | 1.166 | 1,49 | 0.59 | 0.445 | 0.890 | 0.005 | 0.012 |
|  | Training / Pre | 2.44 | 1.220 |  |  |  |  |  |  |
|  | Control / Post | 1.16 | 0.985 |  |  |  |  |  |  |
|  | Training / Post | 0.78 | 1.030 |  |  |  |  |  |  |
| LOC-HIPP | Control / Pre | 1.13 | 0.901 | 1,49 | 2.15 | 0.149 | 0.534 | 0.023 | 0.042 |
|  | Training / Pre | 1.57 | 0.942 |  |  |  |  |  |  |
|  | Control / Post | 1.34 | 0.782 |  |  |  |  |  |  |
|  | Training / Post | -0.84 | 0.818 |  |  |  |  |  |  |
| IFG tri-EC | Control / Pre | 1.25 | 1.283 | 1,49 | 0.07 | 0.786 | 0.911 | <.001 | 0.002 |
|  | Training / Pre | 1.54 | 1.343 |  |  |  |  |  |  |
|  | Control / Post | 0.71 | 0.908 |  |  |  |  |  |  |
|  | Training / Post | 0.34 | 0.950 |  |  |  |  |  |  |
| PRC-IFG tri | Control / Pre | 0.24 | 0.470 | 1,49 | 4.03 | **0.050** | 0.401 | 0.039 | 0.076 |
|  | Training / Pre | 1.31 | 0.491 |  |  |  |  |  |  |
|  | Control / Post | 0.56 | 0.440 |  |  |  |  |  |  |
|  | Training / Post | -0.23 | 0.461 |  |  |  |  |  |  |

*Note.* ANOVA results examining the effect of cognitive training (Group x Time interaction) on connectivity for each individual connection (factors ‘Group’: control, training; between-subject. ‘Time’: pre-, post-training; within-subject). Given are mean values and standard error of the mean (SE), degrees of freedom for the numerator (DFn) and denominator (DFd), p-values (p), and effect size ( $\eta^2_G$ , $\eta^2_p$ ). Within cluster p-FDR correction was applied to the p-values. P-FDR < 0.05 are considered as significant.

**Table S3.** ANOVA cognitive training results for connectivity values in cluster 3.

| Connection | Condition | Mean | SE | DF<sub>nd</sub> | F | p | p-FDR | $\eta^2_g$ | $\eta^2_p$ |
| --- | --- | --- | --- | --- | --- | --- | --- | --- | --- |
| SFG-HPC | Control / Pre | 2.45 | 0.880 | 1,49 | 1.09 | 0.301 | 0.967 | 0.011 | 0.022 |
|  | Training / Pre | 2.80 | 0.921 |  |  |  |  |  |  |
|  | Control / Post | 1.07 | 1.154 |  |  |  |  |  |  |
|  | Training / Post | -0.73 | 1.207 |  |  |  |  |  |  |
| IFG oper-HPC | Control / Pre | 1.61 | 0.729 | 1,49 | 0.01 | 0.921 | 0.967 | <.001 | <.001 |
|  | Training / Pre | 1.29 | 0.762 |  |  |  |  |  |  |
|  | Control / Post | 0.09 | 0.829 |  |  |  |  |  |  |
|  | Training / Post | -0.07 | 0.868 |  |  |  |  |  |  |
| HPC-IFG oper | Control / Pre | 1.86 | 0.897 | 1,49 | 0.01 | 0.925 | 0.967 | <.001 | <.001 |
|  | Training / Pre | 1.44 | 0.938 |  |  |  |  |  |  |
|  | Control / Post | -0.33 | 0.850 |  |  |  |  |  |  |
|  | Training / Post | -0.59 | 0.890 |  |  |  |  |  |  |
| IFG tri-HIPP | Control / Pre | 1.59 | 0.878 | 1,49 | 0 | 0.967 | 0.967 | <.001 | <.001 |
|  | Training / Pre | 1.79 | 0.919 |  |  |  |  |  |  |
|  | Control / Post | -0.29 | 0.784 |  |  |  |  |  |  |
|  | Training / Post | -0.16 | 0.820 |  |  |  |  |  |  |
| HPC-SFG | Control / Pre | 1.54 | 0.669 | 1,49 | 0.12 | 0.733 | 0.967 | <.001 | 0.002 |
|  | Training / Pre | 0.83 | 0.699 |  |  |  |  |  |  |
|  | Control / Post | 0.10 | 0.547 |  |  |  |  |  |  |
|  | Training / Post | -0.99 | 0.572 |  |  |  |  |  |  |
| HPC-IFG tri | Control / Pre | 1.52 | 1.142 | 1,49 | 0.06 | 0.802 | 0.967 | <.001 | 0.001 |
|  | Training / Pre | 1.11 | 1.195 |  |  |  |  |  |  |
|  | Control / Post | -0.31 | 0.859 |  |  |  |  |  |  |
|  | Training / Post | -1.21 | 0.898 |  |  |  |  |  |  |
| IFG tri-IFG oper | Control / Pre | 1.14 | 0.664 | 1,49 | 0.57 | 0.454 | 0.967 | 0.006 | 0.012 |
|  | Training / Pre | 0.54 | 0.695 |  |  |  |  |  |  |
|  | Control / Post | -0.39 | 0.528 |  |  |  |  |  |  |
|  | Training / Post | -0.02 | 0.552 |  |  |  |  |  |  |

*Note.* ANOVA results examining the effect of cognitive training (Group x Time interaction) on connectivity for each individual connection (factors ‘Group’: control, training; between-subject. ‘Time’: pre-, post-training; within-subject). Given are mean values and standard error of the mean (SE), degrees of freedom for the numerator (DFn) and denominator (DFd), p-values (p), and effect size ( $\eta^2_g$ , $\eta^2_p$ ). Within cluster p-FDR correction was applied to the p-values. P-FDR < 0.05 are considered as significant.

#### S2.3 Connectivity change - behavior change

**Figure S3.**
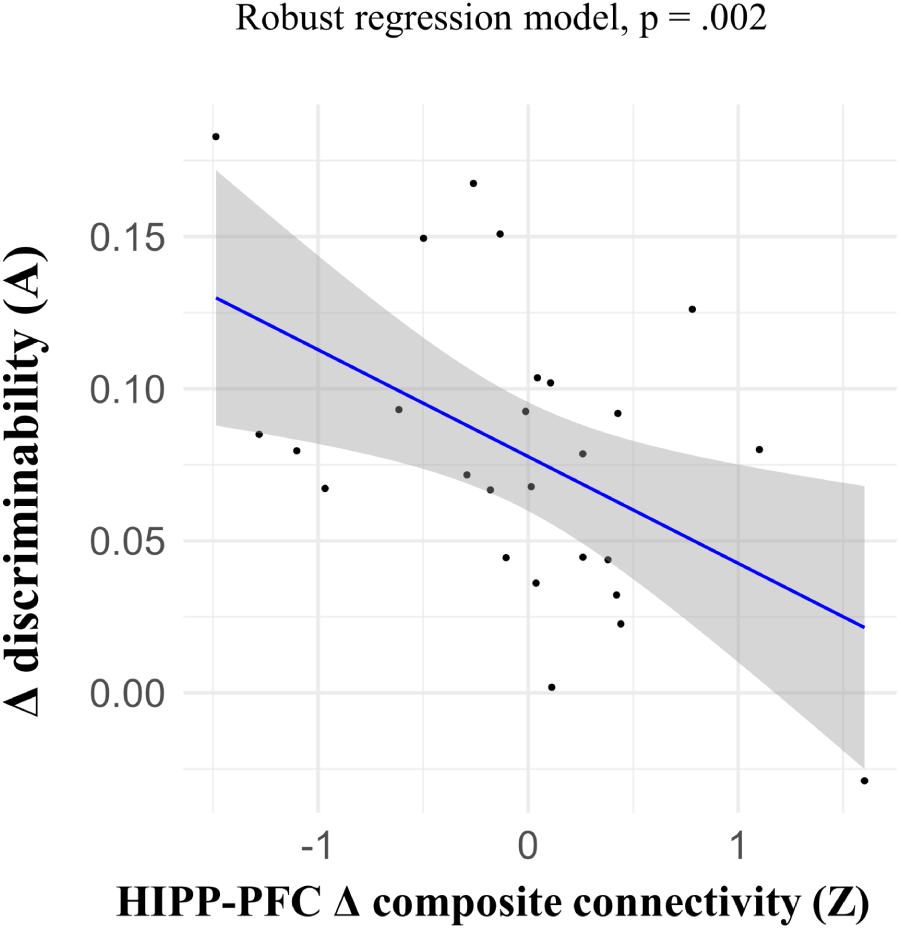
Robust regression: brain connectivity – behavior change. (Δ: post-minus pre-training). Robust linear regression model fitted within the training group. X axis depicts the independent variable: LD connectivity Δ values (gPPI) of a connection (seed ROI – target ROI) or a composite score of a group of connections. Y axis shows the dependent variable (discriminability index A). P-values value for the robust model is shown. **HIPP-PFC**: hippocampal-prefrontal connectivity composite score (Z score; see also section 2.2).

